# Expansion *in situ* genome sequencing links nuclear abnormalities to hotspots of aberrant euchromatin repression

**DOI:** 10.1101/2024.09.24.614614

**Authors:** Ajay S. Labade, Zachary D. Chiang, Caroline Comenho, Paul L. Reginato, Andrew C. Payne, Andrew S. Earl, Rojesh Shrestha, Fabiana M. Duarte, Ehsan Habibi, Ruochi Zhang, George M. Church, Edward S. Boyden, Fei Chen, Jason D. Buenrostro

## Abstract

Microscopy and genomics are both used to characterize cell function, but approaches to connect the two types of information are lacking, particularly at subnuclear resolution. While emerging multiplexed imaging methods can simultaneously localize genomic regions and nuclear proteins, their ability to accurately measure DNA-protein interactions is constrained by the diffraction limit of optical microscopy. Here, we describe expansion in situ genome sequencing (ExIGS), a technology that enables sequencing of genomic DNA and superresolution localization of nuclear proteins in single cells. We applied ExIGS to fibroblast cells derived from an individual with Hutchinson-Gilford progeria syndrome to characterize how variation in nuclear morphology affects spatial chromatin organization. Using this data, we discovered that lamin abnormalities are linked to hotspots of aberrant euchromatin repression that may erode cell identity. Further, we show that lamin abnormalities heterogeneously increase the repressive environment of the nucleus in tissues and aged cells. These results demonstrate that ExIGS may serve as a generalizable platform for connecting nuclear abnormalities to changes in gene regulation across disease contexts.

## Main Text

Since the earliest advancements in microscopy, scientists have identified cells by size, shape, structure, and the appearance of organelles. The morphology of the nucleus is especially important in pathology, as abnormalities in nuclear shape or chromatin texture are routinely used to diagnose cancer and hematologic disorders(*1, 2*). In contrast, with the advent of single-cell sequencing, many now define cell types and states using epigenomic, transcriptomic, and proteomic measurements. Yet for many diseases, the connections between established phenotypic markers and these diverse -omics measurements are not well understood. In recent years, emerging spatial genomics methods have begun to connect microscopy and sequencing modalities, revealing the varied organization of cells within tissues(*3, 4*). While the majority of these approaches focus on measuring transcriptomes at cell resolution, we currently lack methods to connect nuclear abnormalities to changes in genome organization across disease contexts.

The spatial organization of the genome within the nucleus is central to gene regulation(*5, 6*), maintenance of genome integrity(*7*), and the cell cycle(*8*). For cells in the human body to carry out specialized functions, euchromatin localizes to transcriptional hubs, while heterochromatin associates with landmarks like the nuclear lamina for repression and protection from DNA damage(*9*). These functional interactions generally take place at nanometer scale, and disruptions to this precise localization have been implicated in aging and disease phenotypes(*10, 11*). While it seems likely that nuclear abnormalities observed across disease contexts may disrupt spatial genome organization, current approaches are limited in their ability to simultaneously measure both modalities. Optical superresolution and electron microscopy techniques to visualize DNA packing(*12–18*) and epigenetic domains(*19*) at nanoscale reveal general organizing principles, but lack the ability to identify genomic locations. In contrast, emerging whole-genome imaging methods based on DNA FISH measure genomic regions and protein localization(*20–24*), but are constrained in quantifying DNA-protein interactions by the diffraction limit of optical microscopy.

We previously developed *in situ* genome sequencing (IGS)(*25*), a method for sequencing genomic DNA within intact nuclei. IGS combines advantages of microscopy and sequencing approaches – like microscopy, it provides direct 3D localization and is compatible with multiplexed protein imaging; like sequencing, it provides genome-wide measurements. However, the spatial resolution of IGS is subject to the optical diffraction limit, whereas the genomic resolution is constrained by the amount of amplified DNA that fits within the nucleus. To overcome these limitations, we sought to combine IGS with expansion microscopy (ExM)(*26*). In ExM, samples are embedded in a polyacrylate gel and uniformly expanded, allowing for superresolution imaging using a diffraction-limited microscope. While prior methods have used expansion for chromatin imaging(*27, 28*) and *in situ* detection of RNA(*29, 30*), here we develop an approach to simultaneously sequence genomic DNA and image nuclear features at nanoscale resolution.

## Expansion *in situ* genome sequencing

Here, we present expansion *in situ* genome sequencing (ExIGS), a technology for measuring the organization of chromatin within single nuclei at enhanced genomic and spatial resolution. ExIGS integrates and builds upon *in situ* genome sequencing (IGS)(*25*) and expansion microscopy (ExM)(*26*) to enable sequencing of genomic DNA and superresolution immunofluorescence (IF) imaging directly within expanded samples. In developing this protocol, we incorporated strategies from ExM protocols that achieve higher expansion factors(*31, 32*), while also developing new innovations for linking genomic DNA to the gel and optimizing sequencing enzymatics. The resulting workflow for ExIGS consists of three phases: 1) expansion library construction and IF, 2) *in situ* and *ex situ* sequencing, and 3) computational multi-modal integration.

In Phase 1, we link transposed genomic DNA and proteins to a polyacrylate gel for imaging and subsequent sequencing in expanded samples. We first fix cells and perform a mild acidic treatment to make chromatin accessible for whole-genome coverage. We then use Tn5 transposase to integrate sequencing adapters into the genome, creating DNA fragments in their native spatial positions(*33*) (**Fig. 1A, i**). Next, we circularize these fragments by ligating DNA hairpin adapters with unique molecular identifiers (UMIs), followed by immunostaining of proteins (**Fig. 1A, ii**). To preserve the relative spatial locations of genomic DNA and proteins during expansion, we developed an approach where we add DNA oligo hooks complementary to the DNA hairpin containing 5’ acrydite and 3’ amine groups. We then perform a chemical treatment to convert all amines (in both hooks and proteins) to acydite groups that co-polymerize with the gel to anchor both DNA and proteins in place (**Fig. 1A**, iii). We then digest, expand, and re-embed the samples in a secondary gel, followed by superresolution IF imaging (**Fig. 1A**, iv). After IF imaging, we passivate the gel to neutralize its charge and enable enzyme kinetics. Subsequently, we carry out rolling circle amplification (RCA) to generate clonal DNA amplicons for *in situ* sequencing in Phase 2. (**Fig. 1A, v**).

**Figure 1:**
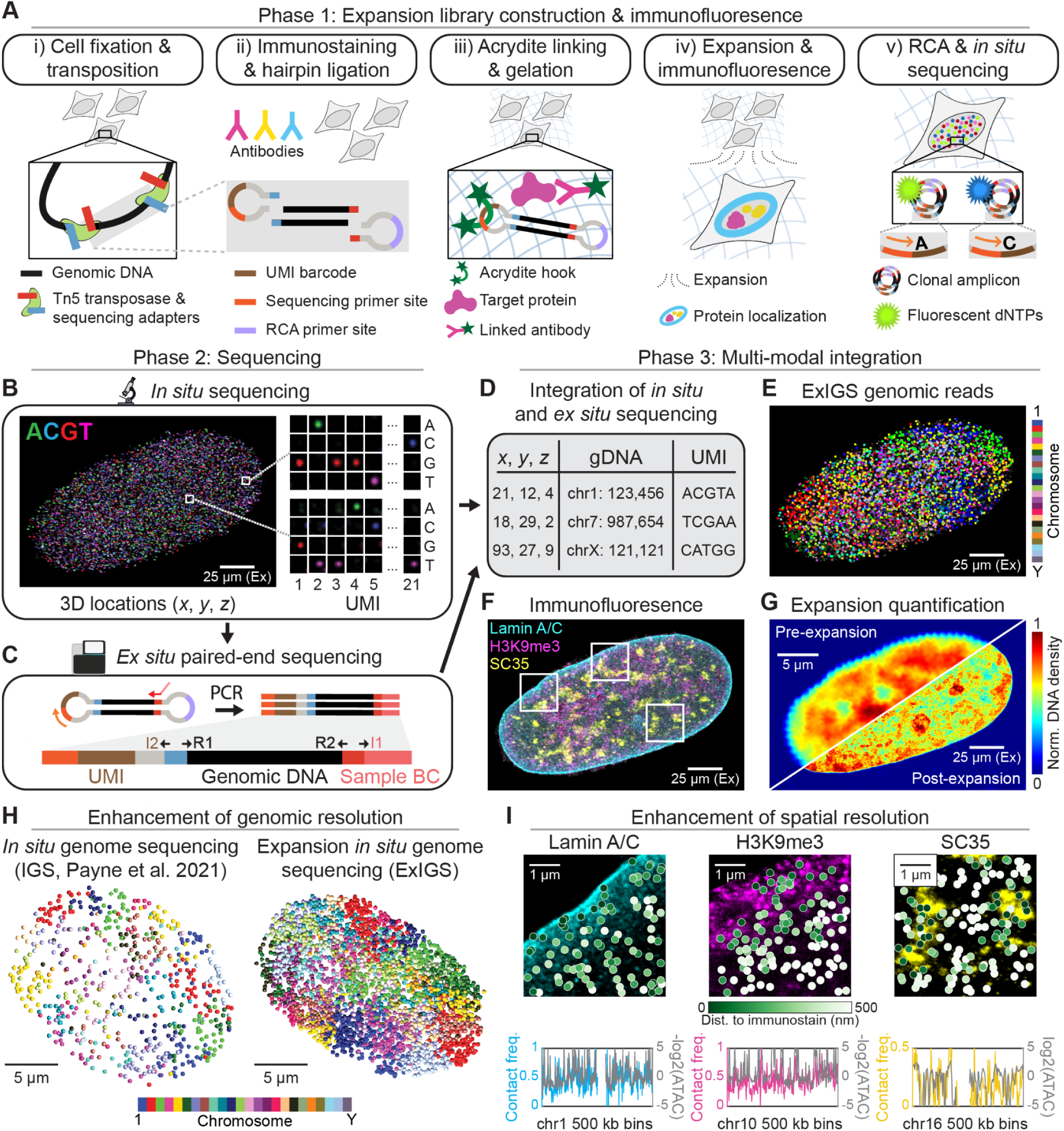
Expansion *in situ* genome sequencing (ExIGS) workflow. (**A**) Workflow for expansion library construction and immunofluorescence (IF). (i) Cell fixation and Tn5 transposition. (ii) Immunostaining and hairpin ligation. (iii) Gel linking and gelation. (iv) Expansion and IF imaging. (v) Rolling circle amplification (RCA) and *in situ* sequencing. UMI, Unique Molecular Identifier; RCA, Rolling Circle Amplification; dNTPs, Deoxyribonucleotide triphosphates. (**B**) Left, one round of *in situ* sequencing-by-synthesis in an expanded IMR-90 nucleus. Right, zoomed in views show representative DNA amplicons with different fluorescent signals for each round of *in situ* sequencing. Up to 21 rounds of sequencing are performed to obtain 3D locations and UMI sequences. (**C**) *In situ* PCR of genomic fragments, followed by *ex situ* paired-end sequencing to obtain genomic DNA and UMI sequences. A sample barcode (BC) is added through a primer handle during PCR for sample multiplexing. (**D**) Table of spatially-resolved genomic reads generated via matching of UMI sequences from (B) and (C). (**E**) Spatially-resolved reads for the IMR-90 fibroblast shown in (B), colored by chromosome. (**F**) Expansion IF imaging of Lamin A/C, H3K9me3, and SC35 for the IMR-90 fibroblast first shown in (B). (**G**) DNA density stain (SYTOX Green) for the IMR-90 fibroblast first shown in (B) before (top left) and after (bottom right) expansion. (**H**) Left, a PGP1 fibroblast with ∼600 spatially-resolved IGS reads(*25*). Right, a skin fibroblast with ∼6,000 spatially-resolved ExIGS reads. Reads colored by chromosome number. (**I**) Top, zoomed in views of the boxes shown in (F) for Lamin A/C, H3K9me3, and SC35 (nuclear speckles). ExIGS points colored by expansion factor-adjusted distance to immunostain. Bottom, plots showing correspondence between immunostain contact frequencies and IMR-90 ATAC-seq for single chromosomes at 500 kb resolution. All images are 3D stacks, but are shown as maximum intensity z-projections for visualization purposes. Scale bars denoted as “Ex” in B, E, F, and G represent observed distances in the expanded sample, while scale bars in H and I represent true expansion factor-adjusted distances.

In Phase 2, we reuse the bi-modal sequencing strategy from IGS(*25*), in which we read out 3D locations and UMI barcodes *in situ*, while acquiring genomic DNA sequences and UMI barcodes “*ex situ*” from an Illumina sequencer. To sequence over 20 bases in an expanded gel, we developed an optimized *in situ* sequencing-by-synthesis (SBS) protocol(*34*) (**Fig. 1B**) with synergistic innovations in gel porosity for increased enzymatic diffusion (**Fig. S1**), a split barcode design to minimize signal decay (**Fig. S2**), and computational correction of spectral bleed-through and molecular phasing (**Fig. S3**). We then perform *in situ* PCR amplification of fragments and *ex situ* paired-end sequencing of genomic DNA and UMIs (**Fig. 1C**).

In Phase 3, we computationally integrate *in situ* and *ex situ* sequencing data, superresolution imaging of nuclear proteins, and expansion factor quantification. We first identify *in situ* 3D amplicons (**Fig. S4**) and match them to *ex situ* paired-end sequencing reads via their UMIs (**Fig. 1D,Methods**), resulting in spatially-resolved genomic reads (**Fig. 1E**). To integrate these reads with the superresolution imaging, we register and segment images for each protein and calculate the nearest distance to every genomic read (**Fig. 1F, Fig. S5**). Lastly, we scale all observed spatial distances between reads and proteins by the expansion factor, which is calculated by computationally aligning a low-resolution image of the sample before and after expansion (**Fig. 1G, Fig. S6**). Collectively, these three phases of the ExIGS workflow enable linking of genomic DNA and proteins, superresolution imaging, bi-modal *s*equencing, and computational data integration in expanded nuclei.

## Validation of ExIGS

We first sought to validate that ExIGS preserves 3D nuclear structure and provides enhanced genomic and spatial resolution compared to methods without expansion. We adapted the gel chemistry from Magnify(*31*), which has been shown to produce a mechanically stable gel with higher expansion factors and minimal distortion. Since the genome is a polymer, we reasoned that transposing genomic fragments prior to expansion would prevent shearing of chromosomes during expansion. By comparing transposed nuclei before and after expansion, we observed that expansion was isotropic, with an expansion factor between 4.5 and 5.5 in all samples (**Fig. S6**). Furthermore, we observed that DNA amplicons were predominantly found within the nucleus and formed a sharp edge at the nuclear boundary.

To assess genomic resolution, we generated ExIGS data from 63 skin fibroblasts (**Table S1**) and compared it to published IGS data of 106 skin fibroblasts(*25*) (PGP1). We observed that ExIGS increased the median number of reads per nucleus from 328 ± 114 in unexpanded fibroblasts to 4,875 ± 1,425 in expanded fibroblasts, an increase of over 10-fold (**Fig. 1H**). As expected, reads from each chromosome occupied distinct spatial territories in the nucleus(*35*) (**Fig. S7**). To further validate that ExIGS preserves 3D genome structure, we clustered reads into homologous chromosomes (**Methods**) and saw that the average pairwise distances between genomic regions strongly resembled Hi-C data (**Fig. S8**). Lastly, we quantified genomic fragment resolution (100-500 bp) and evenness of whole-genome coverage (**Fig. S9**), and found that they were similar to that of IGS.

To evaluate our ability to co-localize DNA and proteins using expansion, we generated ExIGS data from 109 IMR-90 fibroblasts (**Table S2**), a commonly-used cell line in epigenetics and genome structure studies(*36*). This data was paired with expansion IF imaging of Lamin A/C, H3K9me3 (constitutive heterochromatin), and SC35 (nuclear speckles). To quantify spatial resolution, we calculated full width at half maximum (FWHM) values for DNA amplicons and the nuclear lamina with and without expansion, which revealed a substantial improvement for the IF (222 nm with vs. 561 nm without, **Fig. S10**). We also located ExIGS reads relative to each protein stain (**Fig. S5, Methods**); notably, 33.6% of distances were smaller than 200 nm, a lower estimate of the optical diffraction barrier(*37*). To validate the accuracy of protein-DNA contacts, we calculated Lamin A/C, H3K9me3, and SC35 contact frequencies (< 200 nm) for non-overlapping 500 kb genomic bins and compared it to bulk IMR-90 ATAC-seq (**Fig. 1I**). Across the genome, we found that nuclear speckle contact frequency was correlated with chromatin accessibility (ρ = 0.31), whereas nuclear lamina and H3K9me3 contact frequencies were correlated with inaccessibility (ρ = 0.42 and 0.21). Collectively, these analyses demonstrate that expansion can greatly enhance the genomic and spatial resolution of DNA and protein measurements.

## ExIGS connects morphological variation and spatial chromatin organization

One of the main regulatory features of chromatin organization is the nuclear lamina, a meshwork of architectural lamin proteins that resides at the periphery of the nucleus(*38*). Lamin proteins not only maintain the structural integrity of the nucleus(*39*) but also play a functional role in transcriptional silencing(*40, 41*), with prior work demonstrating that localizing genes to the nuclear lamina is sufficient for repression(*42, 43*). While most commonly associated with repressive heterochromatin, there is emerging evidence that lamins may also localize to the nuclear interior(*44, 45*) and interact with euchromatin(*46–48*). However, these interactions have been difficult to characterize with genomic techniques such as ChIP-seq or DamID that do not distinguish between peripheral and internal lamin(*49*).

These difficulties are particularly relevant to the study of laminopathies(*50*), a group of rare genetic disorders that includes Hutchinson-Gilford progeria syndrome (HGPS, henceforth progeria). Progeria causes accelerated aging in children due to a mutation in the lamin A gene that produces an abnormal form of the protein known as progerin(*51*). In cell culture models of progeria, microscopy approaches have demonstrated that the gradual accumulation of progerin results in morphological abnormalities, including internal lamin aggregates(*52*). Relatedly, genomics methods have shown that progeria cells exhibit progressive disruption of epigenetic domains(*53–58*) and genome structure(*54, 59*). Given the unique capabilities of ExIGS to co-localize genomic DNA and nuclear proteins at nanoscale, we decided to use progeria fibroblasts as a model system to directly connect morphological variation to spatial chromatin organization within individual cells.

We performed ExIGS and expansion IF imaging of Lamin A/C on 196 nuclei from a progeria skin fibroblast cell line at passage 19, 22, and 25 (*n* = 64, 134, and 100, **Table S3**). As a control, we used the skin fibroblasts previously examined for validation of genomic resolution (*n* = 63). Inspection of the Lamin A/C images corroborated that the frequency of lamin abnormalities increased with passage(*52*) (**Fig. 2A**). While lamins in control fibroblasts were generally located in a uniform layer at the nuclear periphery (**Fig. 2B**, left); progeria fibroblasts exhibited distorted nuclear shapes, peripheral lamin thickening, and substantial lamin aggregates within the nuclear interior (**Fig. 2C**, left). We next turned to the paired ExIGS sequencing data to examine how these lamin abnormalities affect chromosome organization. Chromosomes in both control and progeria fibroblasts organized into territories (**Fig. 2B, C**, right), but their 3D paths through the nucleus varied based on the lamin environment. In control fibroblasts, chromosomes often traversed both the nuclear interior and periphery (**Fig. 2D**), whereas in progeria fibroblasts, chromosomes more closely followed the contours of the inner lamin topology (**Fig. 2E, Fig. S11**). These combined visualizations highlight the unique observations enabled by combining spatial genomics and superresolution imaging in the same cell.

**Figure 2:**
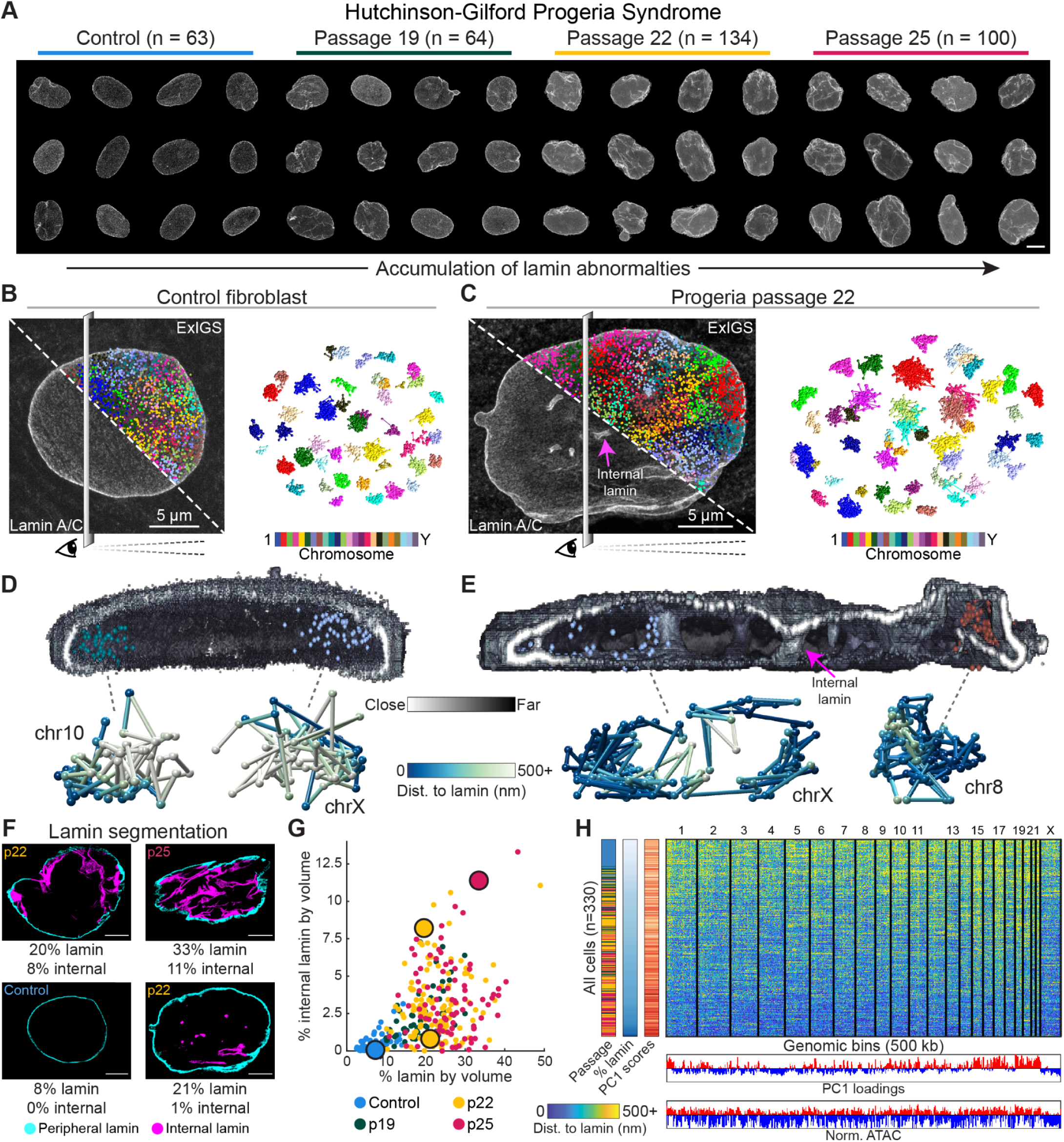
ExIGS connects morphological variation and spatial chromatin organization in progeria fibroblasts. (**A**) Expansion immunofluorescence (IF) images of Lamin A/C for control fibroblasts and progeria fibroblasts from passages 19, 22, and 25. Scale bar in bottom right applies to all nuclei, 5 microns. (**B**) Left, Split visualization of a control fibroblast. Both halves are an expansion IF image of Lamin A/C, top right half also contains overlaid ExIGS reads, colored by chromosome. Dividing plane and eye indicate the perspective shown in (D). Right, Exploded view of homologous chromosomes from the same control fibroblast, with ExIGS reads from each homolog connected by genomic position. (**C**) Same as (B), but for a passage 22 progeria cell. Pink arrow marks an example of an internal lamin structure. (**D**) Top, internal 3D internal of the control fibroblast shown in (B) with two representative chromosomes shown, colored by chromosome number. Lamin is colored by distance from the viewer’s perspective. Bottom, The same two chromosomes shown above, with ExIGS reads connected by genomic position and colored by distance to lamin. (**E**) Same as (D), but for the passage 22 progeria cell shown in (C). Pink arrow marks the same internal lamin structure marked in (C). (**F**) Representative results of the lamin segmentation analysis. Each image is a single *z* plane of a pseudo-fluorescence image, created by splitting the raw fluorescence image into two pseudo-channels using the peripheral and internal lamin segmentation, for visualization purposes only. Scale bars, 5 microns. (**G**) All cells plotted by % lamin by volume and % internal lamin by volume, colored by passage number. Large dots indicate the four cells shown in (F). (**H**) Top, Heatmap showing distance to lamin for 330 single cells by 500 kb genomic bins. Cells are sorted by lamin percentage. Distance to lamin values were averaged across cell reads in a bin and smoothed by 1 Mb in either direction. Bottom, Tracks showing principal component 1 (PC1) loadings and normalized ATAC-seq of control fibroblasts for the same 500 kb genomic bins. All images except (F) are 3D stacks, but are shown as maximum intensity z-projections for visualization purposes.

To move beyond visualization, we developed an image analysis workflow to systematically quantify lamin abnormalities across cells. The workflow classifies nuclear regions occupied by Lamin A/C as either peripheral or internal, then uses these classifications to calculate two metrics: the percentage of each nucleus occupied by 1) total lamin and 2) internal lamin (**Fig. 2F, Fig. S12, Table S4, Methods**). When we applied this workflow to every progeria fibroblast, we observed that both percentages generally increased with cell passage, but exhibited a wide spread, indicative of variation in peripheral lamin thickening and internal lamin (**Fig. 2G**). We also observed that control fibroblasts displayed lower lamin percentages, and validated that we could recapitulate these general trends using non-expansion imaging, albeit at lower spatial resolution (**Fig. S13**). Together, these results demonstrate that ExIGS can capture morphological variation at high resolution across single cells.

We next sought to quantify how lamin abnormalities affect specific regions of the genome. Unsurprisingly, we observed that 98.3% of 500 kb genomic bins were closer on average to lamin in progeria fibroblasts than in controls. We then performed bulk ATAC-seq on control and passage 22 progeria fibroblasts and saw that changes in average distance were correlated with changes in chromatin accessibility (r = 0.31, **Fig. S14**). This suggests that regions with increased proximity to the lamin in progeria fibroblasts generally became more repressed. We then examined these trends at single-cell resolution, ordering all control and progeria fibroblasts by lamin percentage (**Fig. 2H**). While the data was noisy, we saw that many regions localized away from the lamin in control cells, but gradually gained lamin proximity in progeria fibroblasts. When we performed principal component analysis on single-cell lamin distances (**Methods**), we found that per-cell principal component 1 (PC1) scores correlated with lamin percentage (r = 0.46), while PC1 loadings correlated with ATAC-seq tracks of control fibroblasts (r = 0.64). Though most progeria studies focus on loss of heterochromatin(*53–58*), these findings suggest that increased proximity of lamin to euchromatin is a major axis of single-cell variation, with concomitant changes in chromatin accessibility.

## ExIGS reveals local hotspots of disrupted euchromatin organization

One strength of single-cell genomics methods is that they can distinguish between a population of cells in an intermediate state versus a mixture of distinct states. In a similar vein, imaging methods can distinguish between global changes that affect the whole cell versus local changes in subcellular neighborhoods. Given our observations that nuclear abnormalities generally increase lamin-euchromatin proximity in progeria fibroblasts, we hypothesized that this could occur in two distinct ways inside the nucleus: 1) lamin abnormalities cause a global decay of active and inactive chromatin compartmentalization, resulting in aberrant lamin-euchromatin contacts all throughout the nucleus, or 2) lamin abnormalities disrupt euchromatin organization at local hotspots, but leave chromatin compartmentalization in the rest of the nucleus relatively undisturbed.

To distinguish between the “global decay” and “local hotspot” models, we devised a metric to quantify chromatin organization in micron-scale neighborhoods. For each nucleus, we defined a set of spatial neighborhoods containing all ExIGS reads within a 2 micron radius **(Fig. 3A**), then calculated the correlation between distance to lamin (**Fig. 3B**, left) and chromatin activity (**Fig. 3B**, middle, measured via ATAC-seq of control fibroblasts) for each neighborhood. In “organized” neighborhoods with positive correlations, lamin is in closer proximity to heterochromatin, whereas in “disrupted” neighborhoods with negative correlations, lamin is closer to euchromatin (**Fig. 3B**, right). We reasoned that if the global decay model was true, most neighborhoods would have correlations around zero, *i.e.* lamin would have no preference for heterochromatin or euchromatin. In contrast, if the local hotspot model was true, we would observe both strongly organized and strongly disrupted neighborhoods.

**Figure 3:**
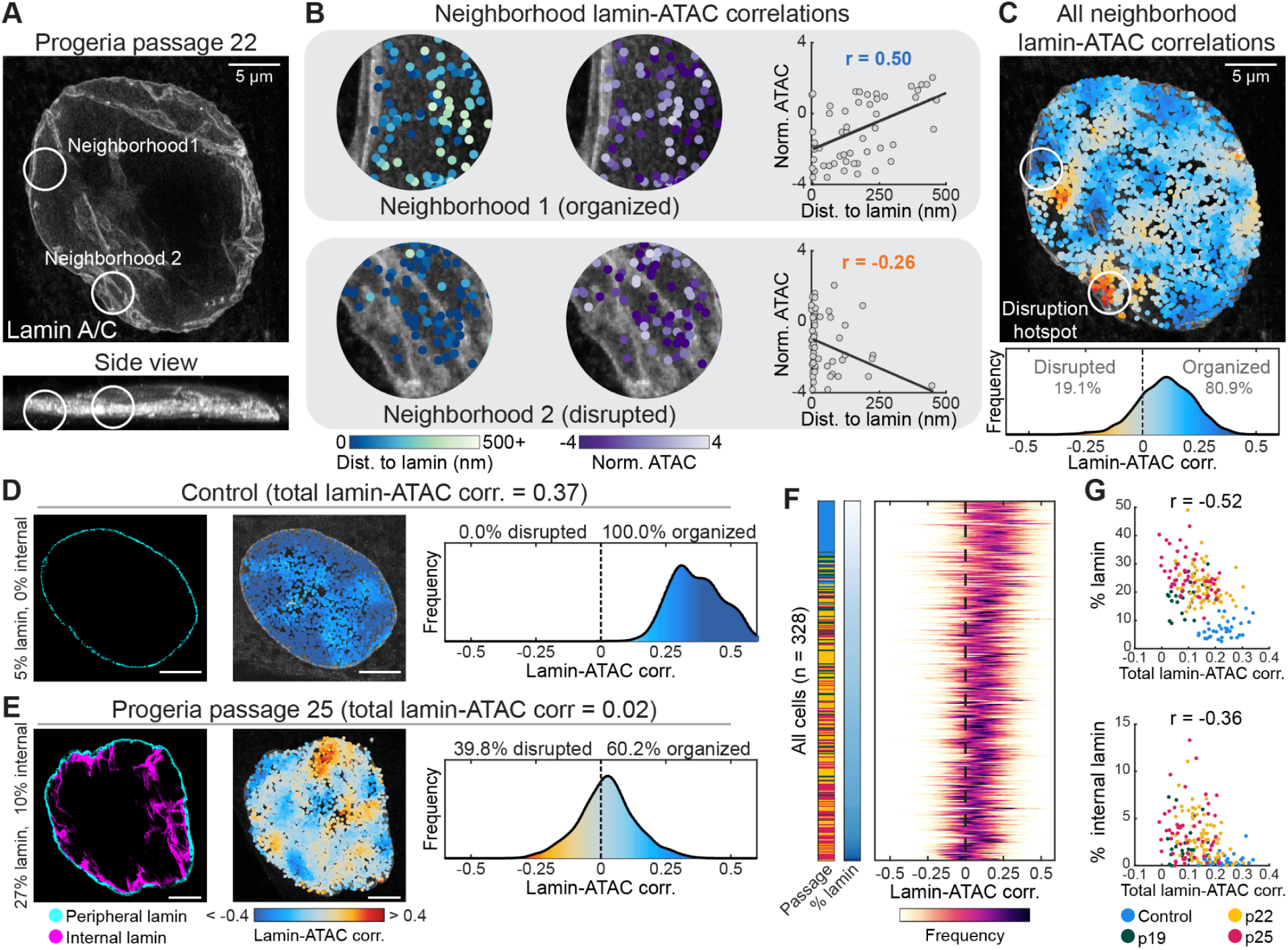
ExIGS reveals local hotspots of disrupted euchromatin organization. (**A**) Top, Expansion immunofluorescence (IF) images of Lamin A/C for a passage 22 progeria fibroblast. Bottom, Side view (maximum intensity projection in *y*) of the same cell. Circles indicate the neighborhoods shown in (B). (**B**) Left, ExIGS reads colored by distance to lamin for the neighborhoods shown in (A). Middle, ExIGS reads colored by normalized 50 kb-binned ATAC-seq of control fibroblasts for the same neighborhoods. Right, Plots of distance to lamin vs. normalized ATAC-seq for the same neighborhoods. Correlation lines and values show the lamin-ATAC correlation for these two neighborhoods. (**C**) Top, Visualization of all neighborhood lamin-ATAC correlations for the progeria fibroblast shown in (A). Bottom, Distribution of all neighborhood lamin-ATAC correlations for the same cell. Dotted line indicates the threshold between “organized” (heterochromatin closer to the lamin) and “disrupted” (euchromatin closer to the lamin). (**D**) Neighborhood lamin-ATAC analysis of a control fibroblast. Left, Image is a single *z* plane of a pseudo-fluorescence image, created by splitting the raw fluorescence image into two pseudo-channels using the peripheral and internal lamin segmentation, for visualization purposes only. Middle, Visualization of all neighborhood lamin-ATAC correlations. Right, Distribution of all neighborhood lamin-ATAC correlations. Scale bars, 5 microns. (**E**) The same as (D), but for a passage 25 progeria fibroblast. (**F**) Distributions of neighborhood lamin-ATAC correlation for all cells, sorted by lamin percentage. Dotted line indicates a correlation of 0. Passage colorbar values are the same as those denoted in (G). (**G**) Top, Mean lamin-ATAC correlations vs. percent lamin by volume for all cells. Bottom, Mean lamin-ATAC correlations vs. percent internal lamin by volume for all cells. Color indicates control or progeria cell passage. All images except (D, left) and (E, left) are 3D stacks, but are shown as maximum intensity z-projections for visualization purposes.

When we calculated all neighborhood correlations for a passage 22 progeria fibroblast, we observed many organized neighborhoods throughout the cell (80.9%), with a few concentrated hotspots of disrupted neighborhoods (**Fig. 3C**). For comparison, we performed the same analysis for a control fibroblast (**Fig. 3D**) and a passage 25 progeria fibroblast with substantial internal lamin (**Fig. 3E**). Remarkably, the control fibroblast had organized neighborhoods throughout (100%), while the passage 25 fibroblasts had fewer organized neighborhoods (61.2%) and a higher incidence of disruption hotspots. When we calculated the distributions for all cells and sorted by lamin percentage, we observed that the distribution mean gradually shifted towards zero (**Fig. 3F**). However, even the most abnormal progeria fibroblasts contained some organized neighborhoods, suggesting that chromatin organization does not globally decay, but rather that disruption is concentrated in local hotspots.

We next sought to quantify the relationship between local disruption hotspots and the accumulation of lamin abnormalities. When we plotted all cells by mean neighborhood correlation and lamin percentage (**Fig. 3G**, top), we observed a strong negative correlation (r = -0.52), corroborating that progeria fibroblasts with more lamin generally exhibit less chromatin organization. We also observed a similar correlation between these mean correlations and percent internal lamin (**Fig. 3G**, bottom), suggesting this disruption might be partially explained by internal lamin aggregates in euchromatic regions of the nucleus. We next asked if disruption hotspots were always located next to internal lamin. Although we found some progeria fibroblasts where this relationship held true, we also observed cases where hotspots were nowhere near internal lamin, as well as instances of internal lamin with no nearby hotspots (**Fig. S15**). This suggests that while the presence of lamin abnormalities is associated with increased frequency of disrupted neighborhoods, they do not directly cause euchromatin localization to the lamin.

Lastly, we sought to characterize if hotspots show a preference for specific genomic regions. We quantified the percentage of reads in each 500 kb genomic bin that fell within a hotspot and observed that hotspots were evenly distributed, with 95% of bins displaying percentages between 3.2% and 9.9% (**Fig. S16)**. Despite the lack of genomic preferences, we hypothesized that random disruptions to euchromatin organization could still adversely affect cell function. To test this, we performed GREAT enrichment analysis(*60*) on all hotspot reads that fell within accessible regions and observed that the most enriched GO cellular components were “actin cytoskeleton”, “anchoring junction”, “focal adhesion”, and other terms associated with cell-to-cell communication and the extracellular matrix (**Table S5**). These results might provide a mechanistic explanation for observations of increased cytoskeletal stiffness and polarity defects in progeria(*61, 62*). More broadly, we posit that lamin abnormalities irregularly disrupt chromatin organization across single cells, and sometimes, these disruptions alter the localization of cell type-specific euchromatin programs.

## Lamin abnormalities are associated with repression and present in tissues and aging

Given our observations of disrupted chromatin organization in progeria fibroblasts, we next sought to determine if these disruptions affect transcription. While lamin proximity is generally sufficient for transcriptional repression(*42, 43*), it is unclear whether lamin abnormalities preserve this function, particularly for internal lamin aggregates within the nucleus(*46–48*). Thus, to quantify transcriptional activity at lamin abnormalities, we performed expansion IF imaging of Lamin A/C and serine-5 phosphorylated RNA polymerase II (Pol II Ser5P, which marks transcriptional initiation) in control fibroblasts (**Fig. 4A**) and progeria fibroblasts from matched passages (19, 22, and 25, **Fig. 4D**). With the nanoscale resolution provided by expansion, we observed a universal depletion of Pol II foci near lamin, whether internal or peripheral (**Fig. 4B, 4E**). In control fibroblasts, the density of Pol II foci was low close to lamin (< 200 nm) and gradually increased with distance (**Fig. 4C**). The same was true of progeria fibroblasts, regardless of whether Pol II foci were near peripheral or internal lamin (**Fig. 4F, Methods**). This indicates that lamin abnormalities are also associated with transcriptional repression, and suggests that the mislocalization of lamin to euchromatin observed in our ExIGS data may be linked to stochastic repression of gene programs important to cell identity.

**Figure 4:**
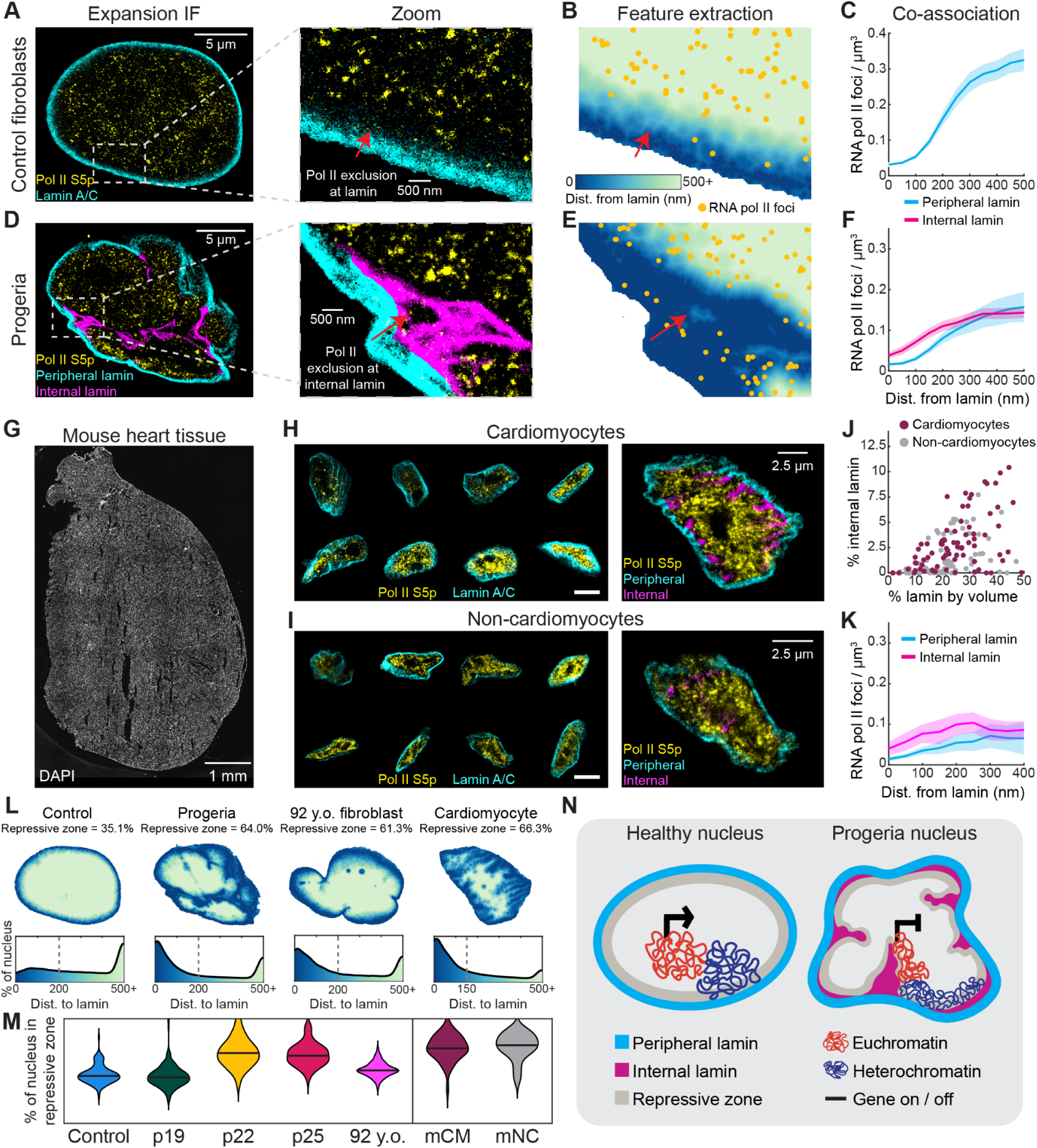
Lamin abnormalities are associated with transcriptional repression and present in tissues and aging. (**A**) Left, Expansion immunofluorescence (IF) imaging of Lamin A/C and Pol II Ser5P. Image is a single *z* plane. Right, Zoomed in view of the boxed region. Arrow denotes a region of low Pol II density near lamin. (**B**) The same region from (A), but with each pixel inside the nucleus colored by minimum distance to the lamin. Yellow dots mark identified Pol II foci. Arrow denotes the same region of low Pol II density. (**C**) Plot showing the Pol II density at increasing distances from the lamin for all control cells (*n* = 118). Upper and lower bounds represent 95% confidence intervals. (**D**) Same as (A), but for a passage 22 progeria fibroblast. Image is a single *z* plane of a pseudo-fluorescence image, created by splitting the raw fluorescence image into two pseudo-channels using the peripheral and internal lamin segmentation, for visualization purposes only. Arrow denotes a region of low Pol II density near internal lamin. (**E**) Same as (B), but for the passage 22 progeria fibroblast shown in (D). (**F**) Same as (C), but for all progeria fibroblasts (*n* = 349). The pink line was calculated using cells with sufficient internal lamin (*n* = 267). (**G**) DNA density (DAPI) image montage of a cryosectioned mouse heart tissue. (**H**) Left, Expansion IF imaging of Lamin A/C (cyan) and Pol II Ser5P (magenta) for select cardiomyocytes (PCM1+). Right, Cardiomyocyte with lamin folds. Image is a pseudo-fluorescence image, created by splitting the raw fluorescence image into two pseudo-channels using the peripheral and internal lamin segmentation, for visualization purposes only. (**I)** Same as (H) but for non-cardiomyocytes (PCM1-). (**J**) All mouse cardiomyocytes and non-cardiomyocytes plotted by % lamin by volume and % internal lamin by volume, as in (2G). (**K**) Same as (F), but for all mouse cardiomyocytes (*n* = 46) and non-cardiomyocytes (*n* =48). (**L**) Top, Representative control fibroblast (left), passage 22 progeria fibroblast (middle left), 92 year-old fibroblast (middle right), and mouse cardiomyocyte (right) with each region in the nuclear volume colored by minimum distance to lamin. Distances shown for a single *z* plane. Bottom, distributions of nuclear volume with respect to lamin. Dotted lines indicate the 200 nm threshold for the repressive zone. (**M**) Violin plot showing the percentage of nuclear volume in the lamin adjacent zone for control fibroblasts, progeria fibroblasts (passages 19, 22, and 25), 92 year-old fibroblasts, mouse cardiomyocytes (mCM) and non-cardiomyocytes (mNC). Repressive zone for mouse cells was defined as being within 150 nm of lamin to account for differing Pol II Ser5P density distributions. Black line denotes the median. (**N**) Schematic of how lamin abnormalities disrupt spatial chromatin organization and transcription in progeria fibroblasts relative to healthy cells. All images are 3D stacks, but select *z* planes are shown for visualization purposes.

We next sought to determine if similar lamin variation exists and affects gene regulation within non-progeria contexts. Though progeria is an extreme phenotype, there is some evidence that lamin abnormalities may be present to varying extents in tissues(*63*) and natural aging(*64, 65*). Since patients with progeria commonly develop cardiovascular disease(*66*), we decided to first focus on lamin variation between and within cell types in the heart. We performed expansion IF imaging of Pol II Ser5P, Lamin A/C, and pericentriolar material 1 (PCM1, a cardiomyocyte marker) on mouse heart tissues (**Fig. 4G**) and observed that cardiomyocytes (PCM1+ cells, **Fig. 4H**, left) and non-cardiomyocytes (PCM1-cells, **Fig. 4I**, left) exhibited diverse nuclear morphologies, including prominent internal lamin folds (**Fig. 4H** and **Fig. 4I**, right). When we systematically quantified these morphologies, we observed similar levels of lamin variation compared to passaged progeria fibroblasts (**Fig. 4J**) and Pol II Ser5P depletion near both peripheral and internal lamin (**Fig. 4K**). Collectively, these results suggest that lamin variation is prevalent in tissues and may modulate transcriptional heterogeneity within a cell type.

We next hypothesized that the abundance and spatial organization of lamin may be an important regulatory mechanism of transcriptional heterogeneity. Given our findings that Pol II foci are depleted in a 200 nm zone surrounding lamin, we leveraged our superresolution imaging to calculate the percentage of each nucleus within this “repressive zone”. In an elliptical control fibroblast, only 35% of the nuclear volume fell within the repressive zone (**Fig. 4L**, left), compared to 64% in a progeria fibroblast (**Fig. 4L**, middle left) and 66% in a mouse cardiomyocyte containing internal lamin folds (with an adjusted 150 nm threshold, **Fig. 4L**, right). Struck by these differences, we decided to test if this mechanism potentially contributes to transcriptional variability in aging(*67, 68*) by imaging a skin fibroblast cell line derived from a 92-year-old donor. We observed substantial lamin abnormalities in these cells (**Fig. S17**), including internal lamin structures and abnormal nuclei that increase repressive volume (**Fig. 4L**, middle right). When we calculated repressive volume percentage for all cells, we observed that old fibroblasts (mean = 51.2%) mostly fell between control (mean = 46.7%) and passage 25 progeria fibroblasts (mean = 65.8%, **Fig. 4M**), suggesting that lamin variability may lead to cellular heterogeneity and transcriptional repression in aging.

Collectively, these results offer a new model of how morphological variability may disrupt healthy gene regulation in aging. In healthy cells, lamins mostly form a uniform layer along the nuclear periphery that contributes to proper silencing of alternative gene programs (**Fig. 4N**, left). However, in models of accelerated aging, lamin gradually accumulates over passaging, which eventually manifests in morphological abnormalities such as abnormal nuclei, thickening of the peripheral lamina, and internal lamin aggregates (**Fig. 4N**, right). As these abnormalities worsen, the environment of the nucleus becomes more repressive, and lamin becomes more likely to disrupt normal chromatin organization. This disruption is not uniformly distributed across the nucleus, but rather occurs in local hotspots. Lastly, because peripheral and internal lamin predominately exclude Pol II, the euchromatin regions within these hotspots are likely repressed, which may heterogeneously affect gene programs important to cell function.

## Discussion

In this work, we describe ExIGS, an approach to unify *in situ* sequencing of genomic DNA and superresolution imaging of proteins. To demonstrate these abilities, we used progeria fibroblasts as a model system to directly connect morphological variability to spatial chromatin disruption in individual cells. We found that lamin abnormalities are linked to hotspots of aberrant euchromatin repression that may disrupt fibroblast activity. Finally, with expansion imaging, we showed that these lamin abnormalities expand the repressive environment of the nucleus, and are prevalent in aging contexts and tissues.

Based on these collective findings, we posit that lamin abundance and organization may be underappreciated factors in transcriptional control and cellular heterogeneity. As we showed in cardiac cells, nuclear morphology is extremely diverse within tissues, and nuclei with increased lamin surface area may exist in more repressed states. Repression is crucial in development as cells differentiate and narrow their range of fates, but has also been observed in aging contexts like senescence, possibly as an adaptation mechanism to prevent aberrant transcription(*69*). This idea is particularly intriguing in light of recent studies suggesting that internal lamin aggregates form as a response to DNA damage(*70*), which may explain how cellular stress triggers repression. Collectively, these observations may even help explain the mechanisms underlying the decay of epigenetic information(*71*) and the increase in transcriptional variability(*72, 73*) reported across aging contexts. Going forward, ExIGS may be applied to study treatments that are thought to slow (or even reverse) progeria and aging phenotypes, such as farnesylation inhibition(*74*) and epigenetic reprogramming(*75*). Beyond aging, we expect that ExIGS will be a generalizable platform for studying gene regulation in any context with diverse nuclear morphologies, such as “stripy” nuclear invaginations in naïve T cells that are predictive of effector function(*76*), or chromatin “herniations” in the nuclear envelope caused by tumor cell migration(*77*).

We anticipate that future iterations of ExIGS will expand measurement capabilities in the molecular, spatial, and temporal dimensions. In the molecular dimension, we expect that ExIGS may be combined with existing expansion-based RNA(*29, 30, 78*) and protein measurements(*79*) via DNA barcodes. In the spatial dimension, integration of ExIGS and iterative expansion protocols that achieve 15 to 20x linear expansion(*80, 81*) may further enhance genomic and spatial resolution. Lastly, in the temporal dimension, it’s possible that our expansion factor quantification algorithm may be adapted to connect live imaging readouts to high-resolution *in situ* measurements(*82, 83*). In summary, we envision that ExIGS will serve as a launching point for a new generation of spatial methods that connect superresolution imaging phenotypes to diverse genomic readouts.

## Acknowledgements

We thank members of the Buenrostro laboratory for critical feedback. We thank S. Ma, S. Mangiamelli, J. Dobkin, A. Sinha, H. Tam, J. Peng, Y. Wang, Z. Dou, P. Griffin, for useful suggestions. We thank the patients and their families for their donations, making this work possible.

## Author contributions

ASL, ZDC, and CC developed the protocol and performed experiments.

ZDC, CC, and JDB performed computational analyses.

PLR and ACP performed proof-of-principle experiments.

GMC, ESB, FC, and JDB supervised proof-of-principle experiments.

ASE, RS, FMD, and EH performed supplementary experiments.

ASE and RZ assisted with computational analyses.

ASL, ZDC, CC, and JDB wrote the manuscript with input from all authors.

JDB secured funding for this work and supervised the study.

## Data and materials availability

All ExIGS data is available in supplementary tables available at Dryad (doi:10.5061/dryad.rr4xgxdhm). Raw sequencing data will be available on the Sequence Read Archive upon publication. All code is available at https://github.com/buenrostrolab/exigs_code.

## Materials and methods

### Cell culture

Human skin fibroblast cell lines (AG08468, AG09602) and Hutchinson-Gilford progeria syndrome cell lines (AG01972) were obtained from the Coriell Institute, while a human lung fibroblast cell line (IMR-90) was procured from ATCC (CCL-186). All cell lines were cultured in Eagle’s Minimum Essential Medium (ATCC #30-2003) supplemented with non-essential amino acids, L-glutamine, 15% fetal bovine serum (FBS, Avantor #89510-186), and 1x Penicillin-Streptomycin (Hyclone #SV30010) at 37°C in a 5% CO_2_. For ExIGS and immunofluorescence assays, approximately 5,000 cells were seeded onto ethanol-sterilized 40 mm coverslips (Bioptechs #40-1313-0319) within silicone gaskets (Grace Bio-Labs #621301) and allowed to adhere for 24 hours.

### Mice

Male wild type C57BL/6 mice were purchased from Jackson Laboratories. Mice were housed in an Association for Assessment and Accreditation of Laboratory Animal Care (AAALAC) approved animal facility at Harvard University. All experimental procedures were performed in accordance with Institutional Animal Care and Use Committee (IACUC) guidelines. During tissue harvest, the heart was collected and freshly embedded in OCT blocks from a 9 week old male mouse.

### *In situ* library construction in expansion gels

#### Cell fixation and permeabilization

Cells (AG08468, AG01972) were washed twice with 1X Phosphate-buffered saline (PBS, Cytiva #SH30256.02) for 5 minutes each, fixed with 4% methanol-free paraformaldehyde (Biotium #22023), in PBS for 10 minutes at room temperature and quenched with 100 mM Tris pH 8 (Corning #46-031-CM) in PBS for 10 minutes with gentle shaking. Following three 5 minute washes with 1X PBS, cells were permeabilized with 0.5% Triton X-100 (Sigma-Aldrich #X100-100ML) in PBS for 10 minutes and washed three times with 1X PBS for 5 minutes. Cells were then treated with 0.1 N HCl for 10 minutes and washed three more times with 1X PBS for 5 minutes each. Next, cells were stained with 1 μg/mL of DAPI (Invitrogen # D1306) in PBS for 10 minutes, washed with 1X PBST (Boston BioProducts #BB-170X) for 5 minutes three times, and a 10 by 10 image montage was taken using a 10x objective and 1x zoom with a 2 μm z-step to identify nuclei prior to expansion.

#### Adapter annealing

Adapter oligos (Ad1 and Ad2) and mosaic end complement oligo (ME) (**Table S6**) were resuspended at 100 μM in nuclease-free water. Adapters were annealed separately with ME (25 μM each of Ad1 and Ad2 adapters, 50 μM ME, 10 mM Tris pH 8, and 50 mM NaCl) using a thermal ramp from 95°C to 25°C over 1 hour in a thermocycler. Annealed adapters were mixed with glycerol (1:1) and stored at -20°C.

#### Tn5 loading and tagmentation

To load the Tn5 enzyme, 1 µL of Tn5 was mixed with 1 µL of annealed adapters and 2 µL of Tn5 loading buffer (50 mM Tris-HCl pH 8, 100 mM NaCl, 0.1 mM EDTA, 0.1% NP-40, 1 mM DTT, and 50% glycerol), and incubated at room temperature for 30 minutes. 2X high salt tagmentation was prepared (66 mM Tris acetate, 132 mM potassium acetate, and 33% DMF, excluding Mg-acetate). We performed two rounds of tagmentation to enhance efficiency. In the first round, cells were incubated in a tagmentation mix (50 µL 2x High Salt Buffer, 5 uL Loaded Tn5, 1 µL PIC, 44 µL nuclease free water) for 1-3 hours at 4°C. The tagmentation reaction was initiated by adding Mg-acetate (Sigma #63052) to a final concentration of 10 mM and incubating cells at 37°C in a humidified box for 1 hour. Following the first tagmentation, cells were washed three times for 10 minutes each with Tn5 wash buffer (50 mM EDTA, 0.01% SDS in 1X PBS) at 45°C, followed by two washes with 1X PBST for 5 minutes each. For the second tagmentation, the cells were pre-incubated with tagmentation mix lacking Mg-acetate for 3 hours at 4°C, followed by the addition of Mg-acetate and incubation overnight at 37°C. After the final tagmentation, cells were washed three times for 10 minutes each with Tn5 wash buffer at 45°C. To de-hybridize blocked ME from DNA fragments, the sample was washed with 40% formamide/2X SSC at 37°C for 15 minutes, followed by two final 5 minute washes with 1X PBST before proceeding with hairpin annealing.

#### Hairpin annealing and hybridization

Hairpin oligos (**Table S6**) were combined at a concentration of 250 nM each in 4x SSC and annealed using a thermocycler programmed with an initial denaturation step at 95°C for 5 minutes, followed by a gradual cooling ramp from 95°C to 20°C at a rate of 1°C per cycle. Cells were incubated with the hairpin mix for 2 hours at 37°C, followed by three 5 minute washes with 1X PBST at room temperature.

#### Gap-fill ligation

Cells were washed with 1X Ampligase buffer (Lucigen #A3210K) at room temperature. Next, the gap-fill ligation mix, containing 1x Ampligase buffer, 50 mM KCl, 2 U/µL Phusion polymerase (NEB #M0530S), 10% formamide, 0.1 U/µL Ampligase (Lucigen #A3210K), and 10 mM dNTPs, was added to the cells. The sample was then incubated for 1 hour on ice with gentle shaking to facilitate diffusion. Subsequently, the cells were incubated at 37°C for 1 hour, followed by 45°C for 1 hour. After incubation, the cells were washed three times with 1X PBST for 5 minutes at room temperature.

#### Immunostaining

After the gap-fill ligation reaction, cells were blocked in 2% BSA (Thermo Scientific #PI37525) in 1X PBST for 1 hour at room temperature. Cells were then stained with primary antibodies diluted 1:200 in 2% BSA /PBST (**Table S7)** overnight at 4°C. After primary antibody staining, cells were washed three times for 10 minutes each with 1X PBST at room temperature. Cells were then stained with secondary antibodies **(Table S7)** diluted 1:200 in 2% BSA in PBST for 1 hour at room temperature. Following incubation, cells were washed three times for 10 minutes each with 1X PBST.

#### Hairpin hook oligo hybridization

To preserve the relative spatial locations of genomic DNA and proteins during expansion, we developed a novel approach involving the use of hook oligos **(Table S6)** that contain a 5’-acrydite and 3’-amine group and are complementary to the DNA hairpins. Following the addition of these oligo hooks, we performed a chemical treatment (Methacrylic acid N-hydroxysuccinimide ester, MA-NHS) to convert all amine groups (present in both the oligo hook and protein) into acrydite groups(*84*). This modification allows the acrydite groups to co-polymerize with the gel network during the expansion gel embedding step. The dual acrydite modifications at both the 5’ and 3’ ends of the oligo hook ensures that all hairpins are securely anchored in place. The hook oligos were hybridized at 2 µM final concentration each in 10% formamide, 4X SSC at 37°C for 3 hours. Cells were then washed twice with 20% Formamide in 2X SSC for 15 minutes each at 37°C, followed by three 5 minute washes with 1X PBST to remove any remaining wash buffer.

#### MA-NHS Treatment

A 1M stock solution of Methacrylic acid N-hydroxysuccinimide ester (MA-NHS, Sigma-Aldrich #730300) was prepared by dissolving 183.16 mg of NHS ester in 1000 μL of anhydrous dimethyl sulfoxide (DMSO) (Thermo Fisher Scientific, catalog number 276855). Aliquots were stored at -20°C in a falcon tube with desiccant (Drierite # 23001) until use. For treatment, an aliquot of 1M MA-NHS ester was thawed and diluted to a 5 mM solution with PBS. Cells were then treated with the 5 mM MA-NHS ester solution for 1 hour at room temperature, followed by three washes with 1X PBST for 10 minutes each.

#### Expansion gel formulation

The Magnify expansion gel formulation was adapted from Klimas et al. 2023 (*31*) and optimized for ExIGS. The optimized formulation provided a dense network of acrylate polymers, ensuring maximum linking efficiency of genomic fragments while maintaining a comparable expansion factor for nanoscale resolution of genomic structures. The final gel composition included 4% N,N-Dimethylacrylamide (Sigma-Aldrich #274135), 6% acrylamide (Sigma-Aldrich #A4058), 34% sodium acrylate (AK Scientific #R624), 0.001% N,N’-Methylenebis(acrylamide) (bis-acrylamide; Sigma-Aldrich #M7279), 1% NaCl (Sigma-Aldrich #S7653), 0.15% ammonium persulfate (APS; Sigma-Aldrich #A3678), and 0.15% N,N,N’,N’-Tetramethylethylenediamine (TEMED; Sigma-Aldrich #T9281) in 1X PBS. The gel stock was prepared and stored in aliquots at -20°C for long-term storage.

#### Sample embedding and expansion

We prepared a gel chamber using a microscope glass slide and strips of scotch tape on both sides, leaving a ∼1 cm wide, ∼60 µm tall gap in the middle. The glass slide was treated with Sigmacote (Sigma #SL2) for 5 minutes to avoid the glass sticking to the gel. The coverslip containing the cells was then inverted onto the treated gel chamber and clamped with paper clamps ensuring the formation of a leak proof chamber around cells. Next, the expansion gel solution (94 µL Magnify gel, 1.5 µL 10% TEMED, 1.5 µL 0.05% 4-Hydroxy-TEMPO (4H, Sigma #176141), 1.5 µL 10% APS and 1.5 µL of H2O) was carefully added to the chamber using a 20 µL pipette tip. The setup was incubated at 4°C for 30 minutes, then transferred to a humidified box and incubated at 37°C for 2 hours to polymerize the gel. The gel was then trimmed to the size of the sample and incubated in digestion buffer (50 mM Tris-HCl pH 8.5, 1 mM EDTA, 0.8 M Guanidine HCl, 0.5% Triton X-100) containing 8 U/mL Proteinase K (NEB #P8107S) for 12-16 hours at room temperature. The gel was allowed to expand by washing it with nuclease-free water for 20 minutes three times.

#### Re-embedding the expanded gel

To stabilize the expanded gel for downstream enzymatics, we re-embed it into a secondary non-expandable acrylamide gel. A glass-bottom 6-well plate was primed for gel adhesion through a treatment of 0.6% Bind-Silane (Sigma-Aldrich #M6514) in 20% Acetic acid 80% Ethanol for 10 minutes, followed by two 5 minute ethanol washes and three water rinses. The expanded gel was cut into smaller pieces, and each piece was transferred to a well of the treated glass-bottom 6-well plate. The gel pieces were re-embedded by immersing them in a cold solution of 2.7% acrylamide (Sigma-Aldrich #A3553), 0.005% APS, and 0.005% TEMED for 10 minutes on ice. Excess acrylamide solution was then removed by aspiration, followed by nitrogen perfusion in a humid airtight container for 10 minutes. The nitrogen perfused container containing the gel was incubated at 37°C for 1 hour for gel polymerization. The re-embedded gel was washed for 5 minutes three times with 1X PBST.

#### Imaging of immunostains

The 6-well glass-bottom plate containing ExIGS samples was transferred to the microscope stage. Each sample well was filled with Illumina imaging buffer and sealed with a transparent PCR plate seal (Bio-Rad #MSB1001). Using Andor Fusion software, multiple fields were acquired to capture immunofluorescence images of expanded cells. Spatial positions and PFS (Perfect Focus System) offsets were saved for each field to facilitate subsequent imaging steps. DAPI imaging was omitted to prevent ultraviolet photocrosslinking. See **Microscopy settings** section for detailed acquisition parameters.

#### Gel Passivation

To neutralize the charge on the gel polymers caused by the sodium acrylate, we performed gel passivation using 1-Ethyl-3-(3-dimethylaminopropyl) carbodiimide (EDC; Sigma-Aldrich #E7750) and N-Hydroxysuccinimide (NHS; Sigma-Aldrich #130672) chemistry in a two-step reaction. Initially, the gel was treated with 150 mM EDC, 150 mM NHS, and 2 M ethanolamine (Sigma-Aldrich #E9508) in 100 mM 2-(N-morpholino)ethanesulfonic acid (MES; Sigma-Aldrich #M1317) buffer (pH 6.5) for 2 hours at room temperature. Following this reaction, the solution was discarded, and the gel was washed with 2 M ethanolamine in 62.5 mM sodium borate (Thermo Scientific Chemicals #AAJ62902AK; pH 8.5) for 40 minutes at room temperature, followed by three 5 minute washes with 1X PBST. Following passivation, the sample was treated with 5 µg/µl SYTOX Green (Thermo Scientific #S7020) and was imaged using a 10x objective to acquire a montage of stained nuclei.

#### Rolling circle amplification

For rolling circle amplification (RCA), a 2 µM locked nucleic acid (LNA) modified RCA primer **(Table S6)** was hybridized in 20% formamide/2X SSC for 3 hours at 37°C. The sample was then washed three times with heated 20% formamide/2x SSC buffer at 37°C for 15 minutes each, followed by three times with 1x PBST for 5 minutes each. Following this, a 16 hour RCA reaction was performed at 30°C using a mixture containing 500 µM dNTP (NEB #N0447S), 50 µM amino-allyl-dUTP (Thermo Scientific #R1091), 1 mM DTT, and 1 U/µL EquiPhi29 in 1X EquiPhi29 buffer (Thermo Scientific #A39392).

#### BS(PEG)9 crosslinking

The gel was washed for 5 minutes three times with 1X PBST and then crosslinked with 5 mM bis-succinimide ester-activated PEG compound (BS(PEG)9, Thermo Scientific #21582) in 1X PBS for 2 hours at room temperature. The crosslinking was quenched by washing the gel with 1 M Tris-HCl (pH 8) for 20 minutes at room temperature, followed by three additional 5 minute washes with 1X PBST.

#### TdT Blocking

Free 3’ OH groups were blocked by incorporating dideoxynucleotides in a terminal transferase (TdT) mixture (200 µM ddNTP mix (AAT Bioquest #17205), 250 µM CoCl2, and 0.4 U/µL terminal transferase in 1X TdT buffer (NEB #M0315S)) for 1 hour at 37°C.

### *In situ* sequencing-by-synthesis

#### Split-barcode design

The DNA hairpins containing UMI sequences were designed such that each 21 base UMI was split into three 6-base barcodes and one 3-base barcode (**Fig. S2, Table S6**). Each barcode was flanked by an upstream primer sequence, enabling sequential readout of individual bases via *in situ* sequencing-by-synthesis. Each barcode was sequenced by hybridizing its corresponding primer, followed by 3-6 rounds of fluorescent dNTP incorporation, imaging, and cleavage. Once all bases of each barcode were sequenced, free 3’-OH groups from sites that were not fully incorporated were blocked to reduce background signal for subsequent rounds of sequencing. The primer of the following primer-UMI pair was then hybridized and the above steps were repeated to sequence all 21 bases of the UMI.

#### Sequencing primer hybridization

A 3 μM LNA modified sequencing primer **(Table S6**) was diluted in 4X SSC buffer and heated at 60°C for 5 minutes to denature secondary DNA structures. The sample was incubated with the heated primer solution at 37°C for 1 hour. The sample was then washed for 10 minutes with PR2 buffer (supplied with Illumina MiSeq kit) three times at room temperature (RT) to remove any unbound or nonspecifically bound primers.

#### Incorporation

Sequencing reagents were extracted from a MiSeq v3 kit (MS-102-3003) reagent cartridge: incorporation mix (position 1), imaging buffer (position 2), and cleavage mix (position 4). These three mixes were aliquoted and stored at -20°C, while the PR2 buffer was stored at 4°C until use. The ExIGS incorporation mix was prepared using 0.5X Illumina V3 incorporation mix, 2.5mM MgCl2, 1x Taq Polymerase buffer (50 mM KCI, 10 mM Tris HCI (pH 8.5 at 25°C), 1.5 mM MgCl, and 0.1% Triton X-100) in PR2 buffer.). The sample was incubated with the ExIGS incorporation mix on ice with gentle shaking for 20 minutes, then transferred to 50°C for 10 minutes. The sample was then returned to the ice block, and fresh ExIGS incorporation mix was added. This process was repeated a total of three times. After the final incorporation, the sample was gently washed for 5 minutes with 1X PBST twice, and then washed with 1X PBST for 10 minutes three times. The sample was subsequently washed with Illumina Imaging Buffer for 10 minutes at room temperature to allow the gel to equilibrate with the buffer. Fresh Illumina Imaging Buffer was then added and the base was imaged.

#### Imaging of sequencing bases

The 6-well glass-bottom plate containing ExIGS samples was placed on the microscope stage. Each well was filled with Illumina imaging buffer and sealed with a transparent PCR plate seal (Bio-Rad #MSB1001). Previously saved XYZ positions and PFS offsets from immunofluorescence imaging were used to relocate the same cells to imaging each base. (See **Microscopy settings** section for detailed acquisition parameters).

#### Cleavage

After imaging, the sample was washed three times for 5 minutes each with 1X PBST. The fluorophores were then cleaved by adding pre-heated Illumina cleavage buffer onto the sample at 50°C on a heating block twice for 10 minutes each. If necessary, photobleaching was performed on the sample after every three bases. To do this, the sample was imaged post-cleavage in cleavage buffer such that each field of view containing cells was given 15 seconds of widefield white light illumination at 100% laser power to minimize background fluorescence. After cleavage (or optional photobleaching), the sample was washed with 1X PBST three times for 5 minutes each, followed by washes of PR2 buffer three times for 5 minutes each at room temperature.

#### TdT Blocking

Once all the barcode bases for the given primer were sequenced, free 3’-OH groups were blocked by incorporating dideoxy nucleotides using a terminal transferase reaction containing 200 µM ddNTP mix (AAT Bioquest #17205), 250 µM CoCl2, and 0.8 U/µL terminal transferase in 1X TdT buffer (NEB #M0315S)). The reaction was incubated for 3 hours or overnight at 37°C. The sample was then washed three times for 10 minutes each with 1X PBST to remove any unincorporated dideoxy nucleotides from the gel before proceeding with the hybridization of the next primer.

### *Ex situ* sequencing

#### DNA extraction

Following in situ sequencing of all UMI bases, in-gel PCR was performed to selectively amplify the UMI and genomic fragment of each DNA amplicon and extract it from the gel. The gel was first washed with water for 10 minutes for a total of five times. Subsequently, the gel was carefully removed from the plate using a razor blade and transferred to a PCR strip tube. The gel weight was measured, and this value was used to adjust the water volume in the PCR mix (1 mg gel = 1 µL water). A 50 µL PCR reaction was prepared containing 1X NEBNext® High-Fidelity PCR Master Mix (NEB #M0541L), 1 µM Ex situ custom forward primer, 1 µM Ex situ custom reverse primer (**Table S6**). This mix was added to the PCR tube containing the gel, and the sample was stored at 4°C overnight to allow for diffusion. The sample was then subjected to an initial 5 PCR cycles, after which the reaction was paused and stored at 4°C. The PCR cycling conditions included an initial denaturation at 98°C for 2 minutes, followed by 5 cycles of 98°C for 15 seconds, 65°C for 30 seconds, and 72°C for 1 minute, with a final extension at 72°C for 5 minutes. To determine the optimal number of additional cycles of amplification, 5 µL of the PCR reaction was used for qPCR (0.5 µM Ex situ custom forward primer, 0.5 µM Ex situ custom reverse primer, 1X NEBNext® High-Fidelity PCR Master Mix, and 1X SYBR Green (Biorad #1708880). The cycle at which the sample reached one-third of the maximum fluorescence (Cycle n) was determined from the qPCR, and the remaining 45 µL PCR reaction was amplified for that many (n) more cycles. Post-PCR, the tube was incubated on a thermomixer at 500 rpm for 3 hours at 4°C to facilitate fragment diffusion out of the gel. Finally, DNA was isolated from the PCR supernatant using a Qiagen column cleanup kit (Qiagen #28006).

#### Paired-end sequencing

Libraries were sequenced on an Illumina Novaseq 6000 platform with 10% phiX. 100bp paired end reads were sequenced with a 8bp i7 to capture the sample barcode and 100bp i5 to capture the in situ UMI sequence.

### Expansion immunofluorescence

#### Preparation of cells

Cells (AG08468, AG09602, and AG01972) were washed twice with 1X PBS for 5 minutes each, fixed with 4% methanol-free paraformaldehyde in PBS for 10 minutes at room temperature (Biotium #22023), and quenched with 100mM Tris pH 8 in PBS for 10 minutes with gentle shaking. Following three washes with 1X PBS, cells were permeabilized with 0.5% Triton X-100 in PBS for 10 minutes and washed again.

#### Preparation of tissue

Fresh frozen heart tissue in OCT was sectioned at 10 µm and placed on an ethanol-sterilized 40 mm coverslip (Bioptechs #40-1313-0319) previously treated with 0.1X poly-L-lysine (Sigma #P8920) for 5 minutes at room temperature. Tissue sections were washed twice with 1X PBS for 5 minutes each, fixed with 4% methanol-free paraformaldehyde in PBS for 10 minutes at room temperature (Biotium #22023), and quenched with 100mM Tris pH 8 in PBS for 10 minutes with gentle shaking. Following three washes with 1X PBS, the tissues were permeabilized with 0.5% Triton X-100 in PBS for 20 minutes and washed again with 1X PBS for 5 minutes three times.

#### Immunostaining

The cells and tissues were blocked in 2% BSA in 1X PBST for 1 hour at room temperature, followed by staining with primary antibodies diluted 1:200 in 2% BSA in PBST (**Table S7)** overnight at 4°C. After primary antibody staining, they were washed three times for 10 minutes each with 1X PBST at room temperature. Cells and tissues were then stained with secondary antibodies **(Table S7)** diluted 1:200 in 2% BSA in PBST for 1 hour at room temperature. Following incubation, they were washed three times for 10 minutes each with 1X PBST.

#### MA-NHS treatment

A 1M stock solution of Methacrylic acid N-hydroxysuccinimide ester (MA-NHS, Sigma-Aldrich #730300) was prepared by dissolving 183.16 mg of NHS ester in 1000 μL of anhydrous dimethyl sulfoxide (DMSO) (Thermo Fisher Scientific #276855). Aliquots were stored at -20°C in a falcon tube with desiccant (Drierite # 23001) until use. For treatment, an aliquot of 1M MA-NHS ester was thawed and diluted to a 5 mM solution with PBS. Cells were then treated with the 5 mM MA-NHS ester solution for 1 hour at room temperature, followed by three washes with 1X PBST for 10 minutes each. Next, the cells and tissue sections were stained with DAPI (1 μg/mL in PBS) for 10 minutes, washed with 1X PBST, and a pre-expansion 10x10 montage was taken using a 10x objective and 1x zoom with a 2-micron Z-step. Nuclei were counted before tagmentation, and pre-expanded nuclei were registered to measure the expansion factor downstream.

#### Expansion gel formulation

The ExIGS Magnify expansion gel formulation adapted from Klimas et al. 2023 (*31*) was used for expansion immunofluorescence. The final gel composition included 4% N,N-Dimethylacrylamide (Sigma-Aldrich #274135), 6% acrylamide (Sigma-Aldrich #A4058), 34% sodium acrylate (AK Scientific #R624), 0.001% N,N’-Methylenebis(acrylamide) (bis-acrylamide; Sigma-Aldrich #M7279), 1% NaCl (Sigma-Aldrich #S7653), 0.15% ammonium persulfate (APS; Sigma-Aldrich #A3678), and 0.15% N,N,N’,N’-Tetramethylethylenediamine (TEMED; Sigma-Aldrich #T9281) in 1X PBS. The gel stock was prepared and stored in aliquots at -20°C for long-term storage.

#### Sample embedding and expansion

We prepared a gel chamber using a microscope glass slide and strips of scotch tape on both sides, leaving a ∼1 cm wide, ∼60 µm tall gap in the middle. The glass slide was treated with Sigmacote (Sigma #SL2) for 5 minutes to avoid the glass sticking to the gel. The coverslip containing the cells was then inverted onto the treated gel chamber and clamped with paper clamps ensuring the formation of a leak proof chamber around cells. Next, the expansion gel solution (94 µL Magnify gel, 1.5 µL 10% TEMED, 1.5 µL 0.05% 4-Hydroxy-TEMPO (4H, Sigma #176141), 1.5 µL 10% APS and 1.5 µL of H2O) was carefully added to the chamber using a 20 µL pipette tip. The setup was incubated at 4°C for 30 minutes, then transferred to a humidified box and incubated at 37°C for 2 hours to polymerize the gel. The gel was then trimmed to the size of the sample and incubated in digestion buffer (50 mM Tris-HCl pH 8.5, 1 mM EDTA, 0.8 M Guanidine HCl, 0.5% Triton X-100) containing 1% of 8 U/mL Proteinase K (NEB #P8107S) for 12-16 hours at room temperature. The gel was allowed to expand by washing the gel with nuclease-free water for 20 minutes three times.

#### Re-embedding of expanded gel

To stabilize the expanded gel for downstream enzymatics, we re-embed it into a secondary non-expandable acrylamide gel. A glass-bottom 6-well plate was primed for gel adhesion through a treatment of 0.6% Bind-Silane (Sigma-Aldrich #M6514) in 20% Acetic acid 80% Ethanol for 10 minutes, followed by two 5 minute ethanol washes and three water rinses. The expanded gel was cut into smaller pieces, and each piece was transferred to a well of the treated glass-bottom 6-well plate. The gel pieces were re-embedded by immersing them in a cold solution of 2.7% acrylamide (Sigma-Aldrich #A3553), 0.005% APS, and 0.005% TEMED for 10 minutes on ice. Excess acrylamide solution was then removed by aspiration, followed by nitrogen perfusion in a humid airtight container for 10 minutes. The nitrogen perfused container containing the gel was incubated at 37°C for 1 hour for gel polymerization. The re-embedded gel was washed for 5 minutes three times with 1X PBST.

#### Microscopy settings

All fluorescence microscopy was performed using an Andor Dragonfly 200 high-speed confocal system coupled to a Nikon Ti2 microscope. Images were acquired with a 40x oil immersion objective (Nikon Plan Fluor, NA 1.30, WD 0.24 mm), a Zyla 4.2 megapixel sCMOS camera with the field of view of 2048x2048 pixels. The system was equipped with PFS (Perfect Focus System) and an ASI MS2000 XY automated stage to enable multi-position imaging. Automated image acquisition parameters were set using Andor Fusion software. For 3D multi-field imaging, a *z*-stack step size of 0.3 µm was used. XYZ positions and PFS offsets were saved for each position within the acquisition protocol to ensure consistent focus and z-stack depth across all fields, thereby reducing acquisition time. The combinations of excitation lasers, dichroic mirrors, and emission filters that were used for imaging are found in **Table S8**.

### Bulk ATAC-seq

#### Adapter annealing and Tn5 loading

Adapter oligos (Ad1 and Ad2) and mosaic end complement oligo (ME) (**Table S6**) were resuspended at 100 μM in nuclease-free water. Adapters were annealed separately with ME (25 μM each of Ad1 and Ad2 adapters, 50 μM ME, 10 mM Tris pH 8, and 50 mM NaCl) using a thermal ramp from 95°C to 25°C over 1 hour in a thermocycler. Annealed adapters were mixed with glycerol (1:1) and stored at -20°C. To load the Tn5 enzyme, 1 µL of Tn5 was mixed with 1 µL of annealed adapters and 2 µL of Tn5 loading buffer (50 mM Tris-HCl pH 8, 100 mM NaCl, 0.1 mM EDTA, 0.1% NP-40, 1 mM DTT, and 50% glycerol), and incubated at room temperature for 30 minutes.

#### Transposition

Cells (AG08468, and AG01972) were washed with 1X PBS, trypsinized (Thermo Scientific #12604013), and centrifuged at 300 × g for 5 minutes, and the pellet was resuspended in cold PBS. For the transposition reaction, 10,000 cells in 5 µL of PBS were mixed with 42.5 µL of 1X High Salt Buffer (33 mM Tris acetate, 66 mM potassium acetate, 10 mM magnesium acetate, 0.1% NP-40, 16% DMF) and 2.5 µL of loaded Tn5 transposase. The reaction was incubated at 37°C for 30 minutes, then the reaction was stopped with 5 µL of 0.5 M EDTA. Transposed DNA was purified using a Qiagen column cleanup kit (Qiagen #28006).

#### Library Preparation

The 15µL of purified DNA was mixed in a PCR reaction containing 1X NEBNext® High-Fidelity 2X PCR Master Mix (NEB #M0541L), 1 µM foward primer, 1 µM reverse primer (**Table S6**). The sample was then subjected to an initial 5 PCR cycles using the following program: 98°C for 2 minutes, followed by 5 cycles of 98°C for 15 seconds, 65°C for 30 seconds, and 72°C for 1 minute, with a final extension at 72°C for 5 minutes, after which the reaction was paused and stored at 4°C. To determine the optimal number of additional cycles offor linear amplification, 5 µL of the PCR reaction was used for qPCR (0.5 µM Ex situ Ad1 primer, 0.5 µM Ad2 primer, 1X NEBNext® High-Fidelity 1X PCR Master Mix, and 1X SYBR Green (Biorad #1708880). The cycle at which the sample reached one-third of the maximum fluorescence (Cycle n) was determined from the qPCR, and the remaining 45 µL PCR reaction was amplified for that many (n) more cycles. DNA was then isolated from the PCR supernatant using a Qiagen column cleanup kit (Qiagen #28006). The library was quantified with a Qubit fluorometer and assessed for quality with a bioanalyzer before sequencing on an Illumina platform, targeting 5 million reads per sample.

### Multi-modal integration

All multi-modal integration steps were implemented using custom MATLAB scripts available from the GitHub repository. Some steps were adapted from *in situ* genome sequencing(*25*), but are described in full for clarity.

#### Field of view image processing

3D nuclei bounds were defined by performing threshold-based segmentation on each field of view from base 1. Nuclei cut off at the edges of fields and cellular debris were excluded from further processing. Fields from each base of sequencing were registered to those from base 1 using normalized cross-correlation to correct for broad xyz offsets between sequencing rounds. Corrected fields were then cropped based on the 3D nuclei bounds, creating five-dimensional nuclei stacks (x by y by z by channels by bases).

#### Nucleus image processing

Cropped nuclei stacks were deconvolved using a high-pass Gaussian filter to improve signal resolution and capped at a maximum pixel intensity to prevent outlier signals from skewing 3D registration. Processed nuclei stacks from each base were then registered to each other using an iterative approach to account for laser-specific spectral aberration. First, nuclei stacks from base 1 were collapsed across the channel dimension using a maximum intensity projection and registered to similarly collapsed stacks from base 2. The resulting transformation was applied to each channel of the (un-collapsed) base 1 stacks, which were then independently re-registered to the collapsed base 2 stack. This same process was then repeated to register stacks from all bases to the registered base1 stack.

#### Correction of sequencing-by-synthesis signals

Nuclei stacks were corrected for spectral bleedthrough and molecular phasing from the optimized sequencing-by-synthesis protocol as follows: First, nuclei stacks for each base were quantile normalized to identify high-confidence amplicons in each channel. These high-confidence amplicons were then used to calculate a linear fit representing the degree of bleedthrough between each permutation of two channels (e.g. bleedthrough was calculated unidirectionally from channel 1 to 2, but also from channel 2 to 1). To correct this bleedthrough, image stacks from the second channel were multiplied by the slope of the fit and subtracted from the first channel, with negative values being capped at 0. Similar fits were calculated for molecular phasing, but for pairs of subsequent bases from the same channel (e.g. a linear fit was calculated between base 1, channel 1 and base 2, channel 1). Following both types of correction, stacks from each channel were subtracted by a minimum intensity value, capped at a maximum intensity value, and divided by the maximum value such that each pixel contains a value between 0 and 1 representing its normalized signal intensity.

#### Quantification of DNA amplicon signals

DNA amplicon centers were identified by applying a 3D peak finding algorithm to registered nuclei stacks from base 1 and base 2, and filtered based on pixel intensity. To quantify the signal intensity of each DNA amplicon, a region-of-interest was defined as a 3 (x) by 3 (y) by 3 (z) pixel volume centered on the amplicon center. Each region was then quantified over all channels and bases of sequencing by summing the fluorescence values of all pixels. The resulting 2D matrix (channels by bases) was normalized such that the sum of squares for each base equaled 1. A purity score for each amplicon was calculated by multiplying the highest values in each base and then performing a negative log transform i.e. -log10(product(max(matrix,1),2)).

#### Alignment of *ex situ* sequencing data

FASTQ files for R1, R2, I1, and I2 were generated using bcl2fastq. Each FASTQ file was then split into chunks of 5M reads using fastp(*85*). The full UMI barcode sequence from I2 was then appended to the read names of R1 and R2 to preserve their association across alignment. Reads were trimmed for adapters, followed by alignment to hg38 with bowtie2(*86*) with –very-sensitive and -k 5 parameters. Aligned reads were then sorted and UMI barcode sequences were moved from the read names to a read group. Chunked BAM files were then re-merged by sample.

#### Deduplication of uniquely aligned, multi-mapping, and unmapped reads

Aligned reads were split into uniquely mapped, multi-mapped, and unmapped subsets using samtools flags(*87*). The first two subsets were further split based on primary alignments, and reads from all subsets were marked as PCR duplicates based on genomic position and UMI sequence using the UMI-tools(*88*) group command with the --edit-distance-threshold 2 parameter. Subsets were then merged based on mapping quality, with multi-mapped and unmapped reads being collapsed into a single entry per UMI. The resulting list of UMIs was filtered for index swapping based on the frequency of each unique UMI-genomic location combination.

#### Annotation of repetitive DNA elements

Both uniquely mapped and multi-mapped reads were annotated based on their overlap with repetitive DNA elements from the RepeatMasker database(*89*) using bedtools(*90*). For multi-mapped reads, overlap was calculated for all potential alignments and collapsed into a single entry. Each overlapping read was annotated at the repeat name, family, and class levels.

#### Matching of *in situ* and *ex situ* UMIs

Spatially-resolved ExIGS reads were generated by matching the normalized UMI probability matrices from *in situ* sequencing images to the UMI sequences read out via *ex situ* sequencing. A consensus *in situ* UMI sequence was generated for each 3D amplicon location by selecting the imaging channel with the highest probability per base. For each consensus *in situ* UMI, a list of *ex situ* UMIs with Hamming distances less than 4 was generated. A match score was calculated for each *ex situ* UMI by multiplying the corresponding probabilities from the *in situ* probability matrix and performing a negative log transform. The *ex situ* UMI with the lowest match score per 3D location was selected to create ExIGS reads with matched spatial and genomic positions. To match each nucleus to its corresponding *ex situ* sequencing sample, the first 1000 3D amplicon locations were processed and the sample with the most matches was selected. ExIGS reads were filtered using both purity scores and match scores. The threshold was adjusted for each cell by calculating the number of informative bases and setting the y-intercept to be 10 - 0.75*(non-informative bases). Cells with less than 14 informative bases were filtered out.

#### Quantification of expansion factor

The 10x or 20x image montage of the coverslip prior to expansion was projected into 2D via maximum intensity *z*-projection and split into tiles for nuclei segmentation. An Ilastik(*91*) model was trained per montage and applied to all tiles. To get the locations of nuclei following expansion, either a 10x image montage of the gel was taken or an equivalent montage was reconstructed by stitching together all 40x fields-of-view. Nuclei in the post-expansion montage were also projected into 2D and segmented using Ilastik.

To register the pre- and post-expansion montages, algorithms were developed to calculate unique “positional fingerprints”, which leverage that the relative spatial locations of nuclei are preserved during expansion. For each nucleus, its nearest neighbor by Euclidean distance was found and normalized to an angle of 0 degrees and a distance of 1. Next, the positions of all other neighboring nuclei were calculated relative to this normalized angle and distance. Nuclear fingerprints were then compared for all pairs of pre- and post-expansion nuclei, with neighbors within 15 degrees and distances of 0.8 - 1.2 being considered as matches. To minimize computational resources, matching for each pair of nuclei stopped after 25 non-matches were found. After matches for all pairs were counted, the pairs with the highest numbers of matches were visualized and used to calculate an affine transform between the pre- and post-expansion montages.

The resulting affine transform was used to compare images of individual nuclei before and after expansion. For registration, 5x was used as an initial estimate of the expansion factor. Images were registered in an iterative process, first with a rigid transformation that considers only translation and rotation, then with a similarity transformation that considers scale as well. The correlation of the resulting images was used to assess registration quality. For nuclei with sufficient registration quality, the scale factor was derived from the similarity transform matrix and multiplied by 5 to calculate the expansion factor. The expansion factors for nuclei within a sample were very consistent, so the mean expansion factor was applied to all nuclei.

#### Quantification of expanded resolution

To quantify the feature width of DNA amplicons and the nuclear lamina with and without expansion, pixel intensity was measured across a 1 micron line scan. For the quantification of nuclear lamina feature width with expansion, ExIGS immunofluorescence of Lamin A/C in a human skin fibroblast was used, whereas for without expansion, IGS immunofluorescence of Lamin B in mouse early embryos was used. To calculate full width at half max (FWHM) for each feature, first, the position corresponding to the max value in the line scan was found. Then the nearest local minima on either side of this position were identified, and the lower value was selected to calculate the halfmax value using the following equation: halfmax_value = (max_value - lower_min)/2. The positions of the first and last positions with values greater than the halfmax value were then found and resolved further using linear interpolation. Finally, the distance between these interpolated points was taken as the FWHM.

#### Processing of expansion immunofluorescence imaging

Paired expansion immunofluorescence images were registered to 3D amplicon locations using an interactive registration approach. 3D image stacks were first broadly aligned to the first base of *in situ* sequencing using cross-correlation. Image stacks were then cropped based on nuclei bounds and then re-registered using image filling strategies that impute the overall shape of nuclei. Laser-specific spectral aberrations were corrected using fluorescent beads or background signals present in all channels. For each non-Lamin A/C protein immunostain, an intensity-based segmentation threshold was set for each sample based on visual inspection.

#### Segmentation of peripheral and internal lamin

To define an overall region-of-interest for lamin segmentation, a convex hull was generated based on base 1 of *in situ* sequencing. Two pixel intensity-based thresholds were then set – the first to define the locations of Lamin A/C, the second to define the bounds of the nuclear volume. Each pixel defined as lamin by the first threshold was then labeled as either peripheral or internal lamin based on the minimum distance to the edge of the nuclear volume. Pixels with an expansion-adjusted distance over 600 nm were labeled as internal lamin; all other lamin pixels were labeled as peripheral lamin. For validation in unexpanded progeria fibroblasts, a threshold of 1 micron from the edge of the nuclear volume was used to label internal lamin. ExIGS reads in each nuclei were annotated by their minimum distance to both labels. The number of pixels with each label was then summed and used to calculate the percentage of each nuclear volume occupied by total lamin and internal lamin. The volume and axes lengths of each nucleus were also saved as per-cell statistics. ExIGS reads more than 200 nm outside the nuclear volume were marked and excluded from downstream analysis.

#### Quantification of Pol II Ser5P density and repressive lamin zones

These quantifications depend on the Lamin A/C segmentation, the nuclear volume segmentation, and the peripheral/internal lamin labels described above. For each pixel in the nuclear volume, a minimum distance to the lamin was calculated in voxel space and adjusted by expansion factor. Pixels were binned into 50 nm increments and labeled by whether they were closer to peripheral or internal lamin. Pol II Ser5P foci were identified by applying a high-pass Gaussian filter to deconvolve raw fluorescence images, which were used as input to a 3D peak calling algorithm. For each cell, the density of Pol II Ser5P foci was calculated at various distances from both peripheral and internal lamin. Nuclei with less than 5000 internal lamin pixels were excluded from the internal lamin distribution. The mean and 95% confidence intervals across all cells were plotted for each distance increment to visualize the exclusion of Pol II Ser5P foci near both peripheral and internal lamin.

#### Assignment of ExIGS reads to homologous chromosome clusters

Nuclei with more than two spatial clusters per chromosome were removed by visual inspection. The following process was performed for each autosome and chromosome X for every cell: First, a pairwise distance matrix was generated for all ExIGS reads aligned to a specific chromosome. The distances for each read were sorted, and reads with a 20th percentile distance less than 20% of the max pairwise distance were filtered out. Next, DBSCAN clustering was performed with a neighborhood search radius of 10 and a minimum neighbor threshold of 5, and points not belonging to a cluster were also filtered. K-medoids clustering was then performed with k = 2, these basic spatial clusters were used as the input to the following block swapping algorithm.

To deconvolve intermingling homologs, an algorithm was developed to weigh genomic position in addition to spatial location. First, using the spatial cluster assignments from above, a distance was calculated for each cluster by sorting ExIGS reads by genomic position and calculating the length of the path in 3D space. The algorithm then attempted to swap every possible consecutive block of 5 reads or less to the other cluster. If the resulting clusters lowered the combined length of the two homologs, the swap was kept. The algorithm stopped when there were no more swaps that would lower the combined distance. Lastly, reads that contribute disproportionately to the path length were filtered out. The remaining reads were labeled as belonging to high-confidence clusters for downstream analysis.

### Genomic analyses

#### Comparison to Hi-C data

Hi-C data of IMR-90 fibroblasts was downloaded from GSE63525(*92*) and lifted over to hg38 using HiCLift(*93*). Lifted over contacts were then converted to .hic format, normalized using Knight-Ruiz matrix balancing, and outputted as a contact frequency matrix at 1 Mb resolution using Juicer(*94*). To compare ExIGS to Hi-C, ExIGS reads were assigned to non-overlapping 1 Mb genomic bins. For every high-confidence cluster in each IMR-90 cell, a pairwise distance matrix was constructed for all reads in the cluster. These distances were used to construct a mean spatial distance matrix per chromosome, where the value of each element represents the average distance between a pair of genomic bins across all of the IMR-90 ExIGS data. For each chromosome, a Pearson correlation was calculated between the values of this mean spatial distance matrix and the corresponding values of the normalized Hi-C contact frequency matrix.

#### Processing of bulk ATAC-seq data

For bulk ATAC-seq of control and progeria passage 21 fibroblasts, FASTQ files were generated using bcl2fastq. Reads were trimmed for adapters, followed by alignment to hg38 with bowtie2(*86*). Aligned reads were then sorted and filtered using samtools, followed by deduplication of PCR duplicates. For various integrations with ExIGS data, reads were partitioned into non-overlapping 50 and 500 kb genomic bins spanning hg38. For bulk ATAC-seq of IMR-90, aligned BAM files were downloaded from ENCSR200OML(*95*)

#### ExIGS contact frequency at nuclear landmarks

ExIGS reads from IMR-90 fibroblasts were partitioned into non-overlapping 500 kb bins spanning hg38. For each bin, the number of reads contacting (< 200 nm away) each immunostain were divided by the total number of reads in the bin, yielding a contact frequency between 0 and 1. Spearman correlations were calculated using the 500 kb contact frequencies for each immunostain and the log2 transformed binned ATAC-seq values from IMR-90. For Fig. 1G, the sign of the ATAC-seq values were flipped for the purpose of visualizing concordance with Lamin A/C and H3K9me3, which are associated with repressed heterochromatin.

#### Comparison of ATAC-seq and distance to lamin changes in progeria fibroblasts

The normalized 500 kb binned ATAC-seq values of passage 21 progeria fibroblasts were divided by that of control fibroblasts and log2 transformed. The mean spatial distance of each 500 kb bin to lamin was calculated for all progeria fibroblasts and the corresponding mean distances of control fibroblasts was subtracted. The Pearson correlation between the log2 transformed ATAC-seq values and the differences in distances to lamin was calculated.

#### Neighborhood lamin-ATAC scores

A spatial neighborhood was defined for each ExIGS read by identifying all neighboring reads within a 200 nm radius. All ExIGS reads were annotated using the log2 transformed ATAC-seq value of the surrounding 50 kb genomic bin. For each neighborhood, a Pearson correlation was calculated using these normalized ATAC-seq values and the minimum distance of each ExIGS read to the lamin. Correlations above 0 were labeled as “organized”, whereas those below zero were labeled as “disrupted”. To calculate a total lamin-ATAC correlation per cell (**Fig. 3G**), the neighborhood definitions were ignored, and a single correlation was calculated for all ExIGS reads within the cell.

#### Characterization of disruption hotspots

Neighborhoods with a lamin-ATAC correlation under -0.1 were denoted as “disruption hotspots”. To quantify hotspot frequency across the genome (**Fig. S16**), all ExIGS reads were partitioned into non-overlapping 500 kb genomic bins, and the number of reads in a hotspot were divided by the total number of reads in each bin. Bins with less than 10 reads were excluded. For the GO cellular components enrichment analysis, reads in hotspots were inputted to GREAT(*60*)

## List of Supplementary Figures

**Fig. S1:**
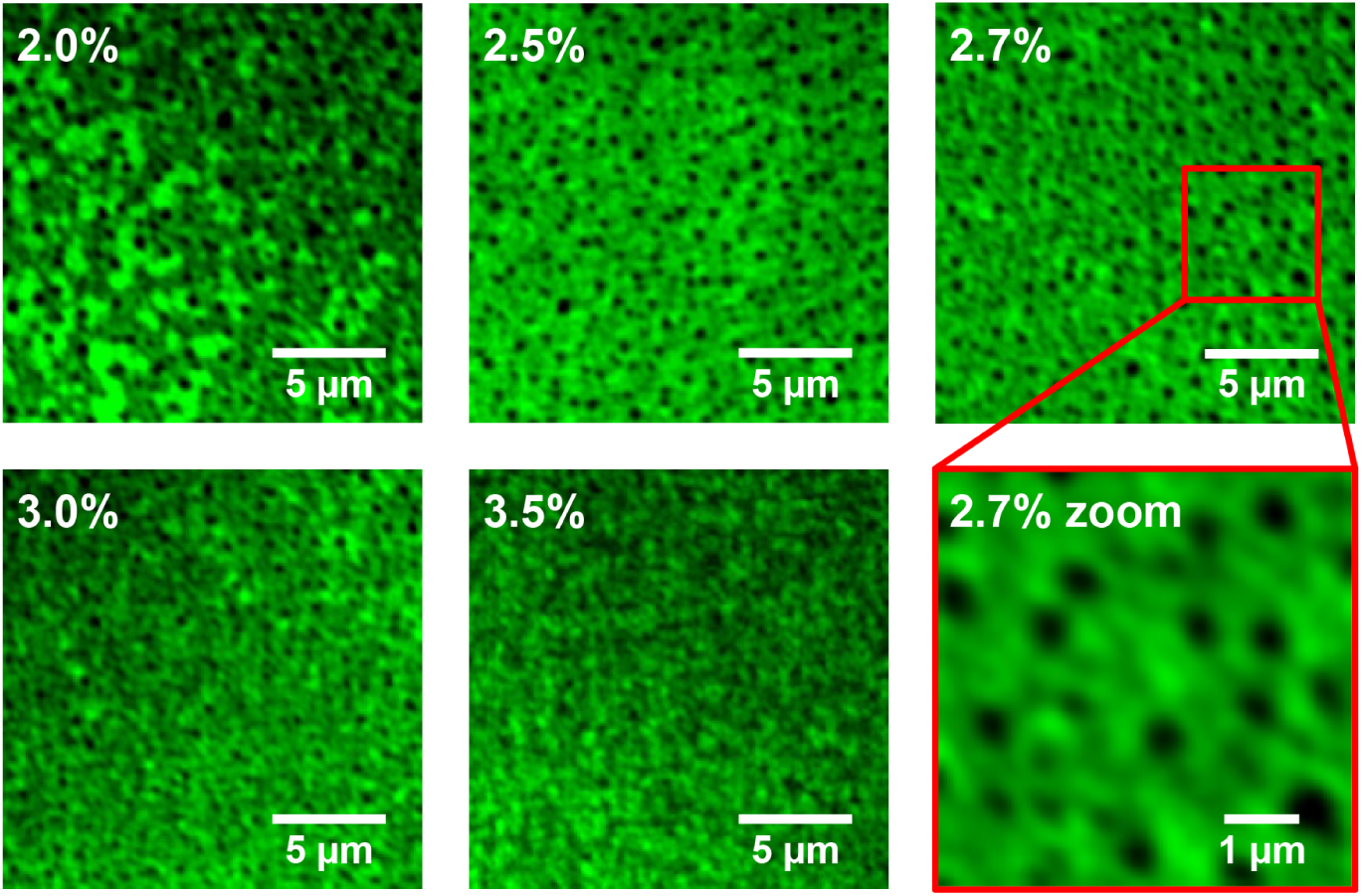
Optimization of re-embedding gel composition. Images of secondary re-embedding gels made using different Acry:Bis ratios, visualized with an acrydite-modified fluorescent moiety. Higher acrylamide percentages form more dense gels with smaller pores. We found that low acrylamide gels (2-2.5%) fail to completely polymerase, and higher acrylamide gels (3-3.5%) have small pore size which disrupt diffusion of enzymes into the gel. We found that 2.7% acrylamide gels polymerize and retain large enough pores to facilitate enzyme diffusion into the gel.

**Fig. S2:**
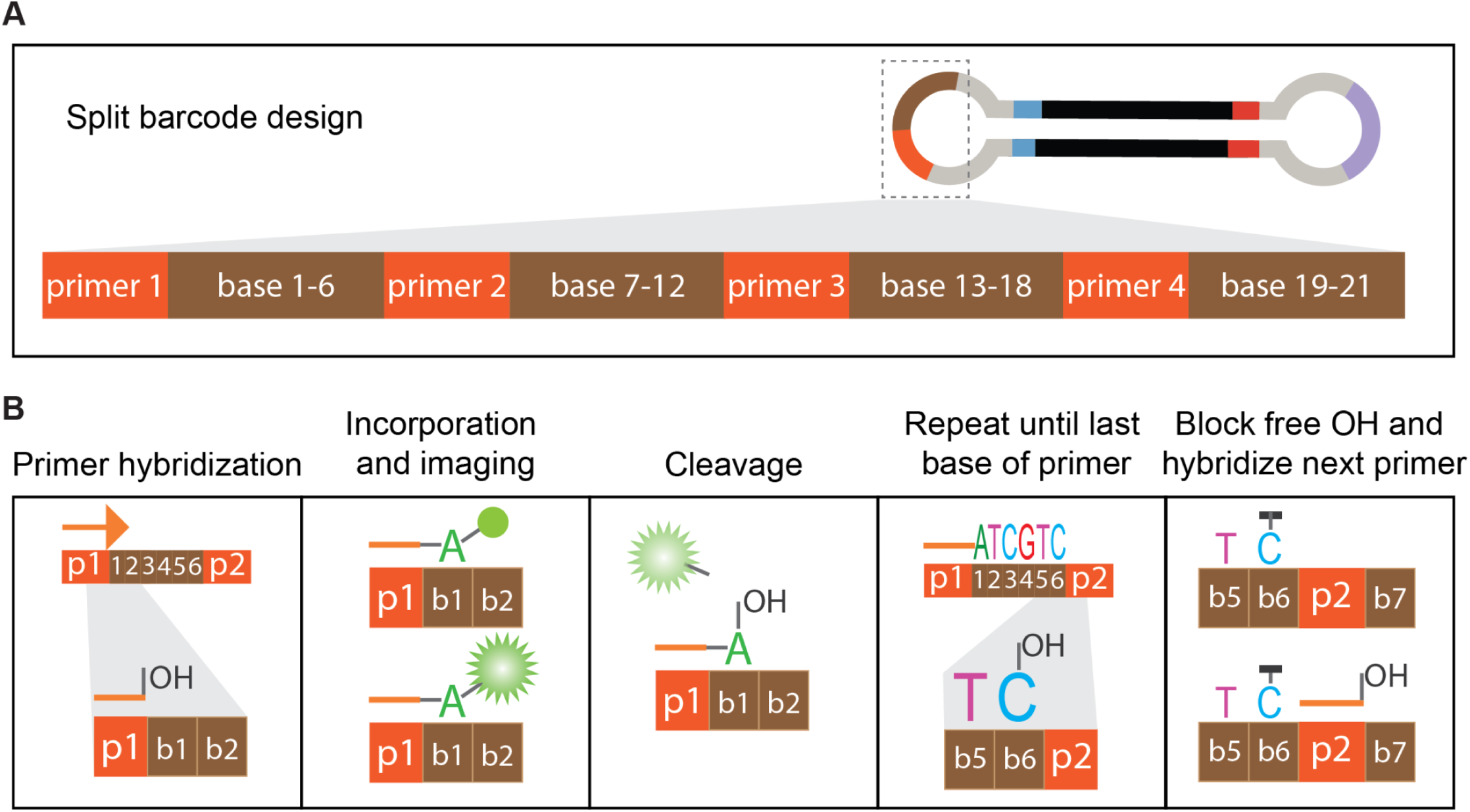
Split barcode design for sequencing-by-synthesis. (A) Schematic of the split barcode design used for *in situ* sequencing-by-synthesis. Including multiple primers and performing a blocking step before each primer “resets” the gradual decay of fluorescent signals caused by signal phasing. (B) Workflow for *in situ* sequencing-by-synthesis using the split barcode design shown in (A). Primer hybridization is followed by repeated dNTP incorporation, imaging, and cleavage. Once all of the bases of a primer is read out, free -OH groups are blocked and the process is repeated for each primer.

**Fig. S3:**
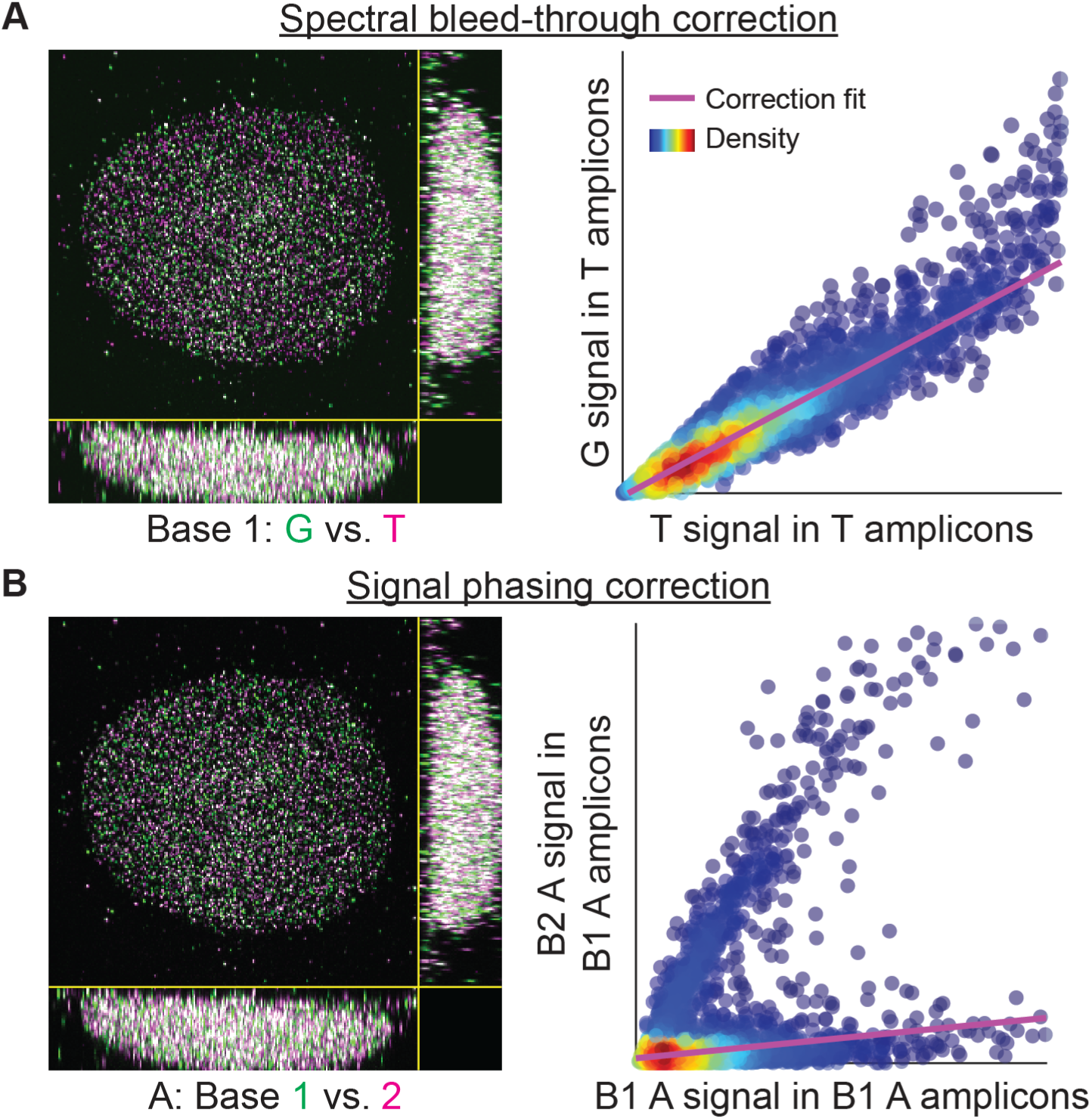
Correction of sequencing-by-synthesis fluorescence signal. (A) Computational correction of spectral bleed-through. For each base and channel, a set of high-confidence DNA amplicons is defined, and fluorescent signal is measured at these locations for all other bases (e.g. we define a set of high-confidence T amplicons and measure the signal at these locations in A, C, and G). These signals are used to calculate a fit that represents the degree of bleed-through between each pair of channels, and are applied to correct the original images. (B) Same as (A), but for signal phasing. For each base and channel, a set of high-confidence DNA amplicons is defined, and fluorescent signal is measured at these locations in the next base (e.g. we defined a set of high-confidence A amplicons in base 1 and measure the A signal at these locations in base 2). These signals are used to calculate a fit that represents the degree of bleed-through between subsequent bases, and are applied to correct the original images.

**Fig. S4:**
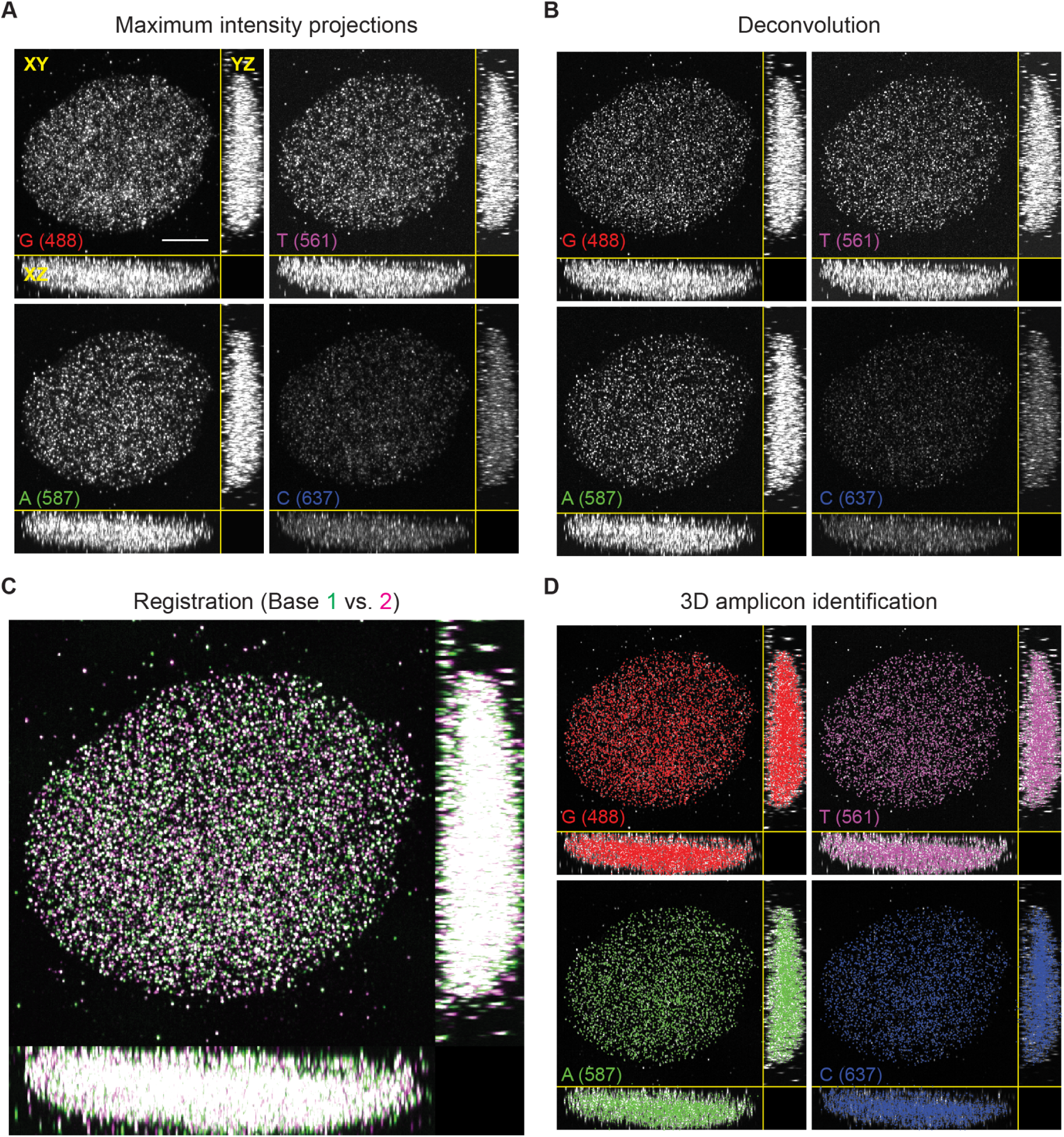
Registration and identification of 3D amplicon positions. (A) Images from each channel of *in situ* sequencing for base 1 of a skin fibroblast. Each image shows maximum intensity projections in *x, y*, and *z*. (B) The same images as (A), but following Gaussian high-pass filtering. (C) The nucleus first shown in (A) after registration of base 2 images to base 1 images. Green represents signal only in base 1, magenta represents signal only in base 2, and white represents signal in both. (D) The same images as (B), but with overlaid amplicon locations identified via 3D peak calling.

**Fig. S5:**
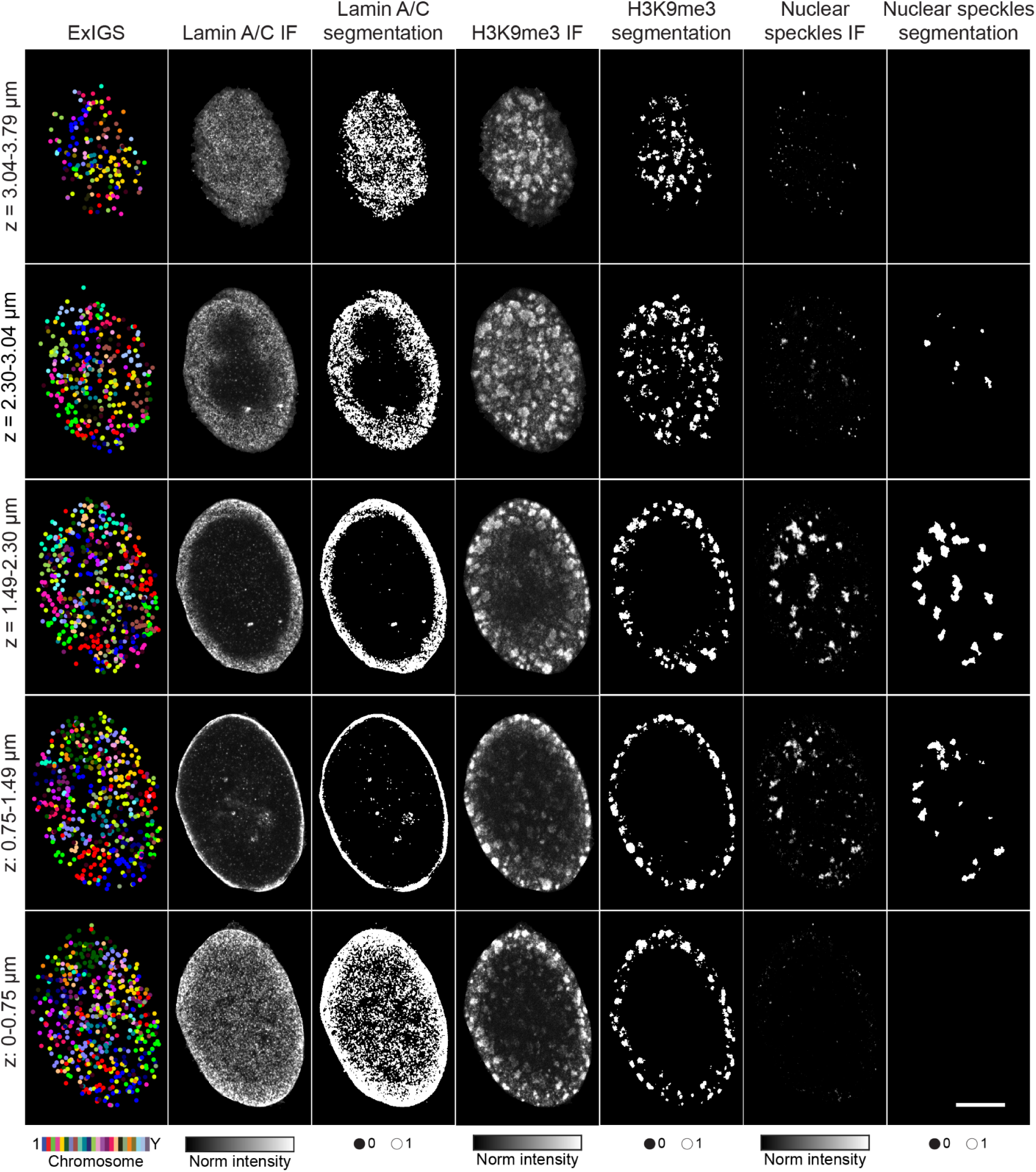
Expansion immunofluorescence imaging and segmentation. Image stacks showing either expansion immunofluorescence imaging of targeted nuclear proteins (Lamin A/C, H3K9me3, and nuclear speckles) and their segmentation at various 3D slices of an IMR-90 fibroblast. Scale bar in bottom right applies to all images, 5 microns.

**Fig. S6:**
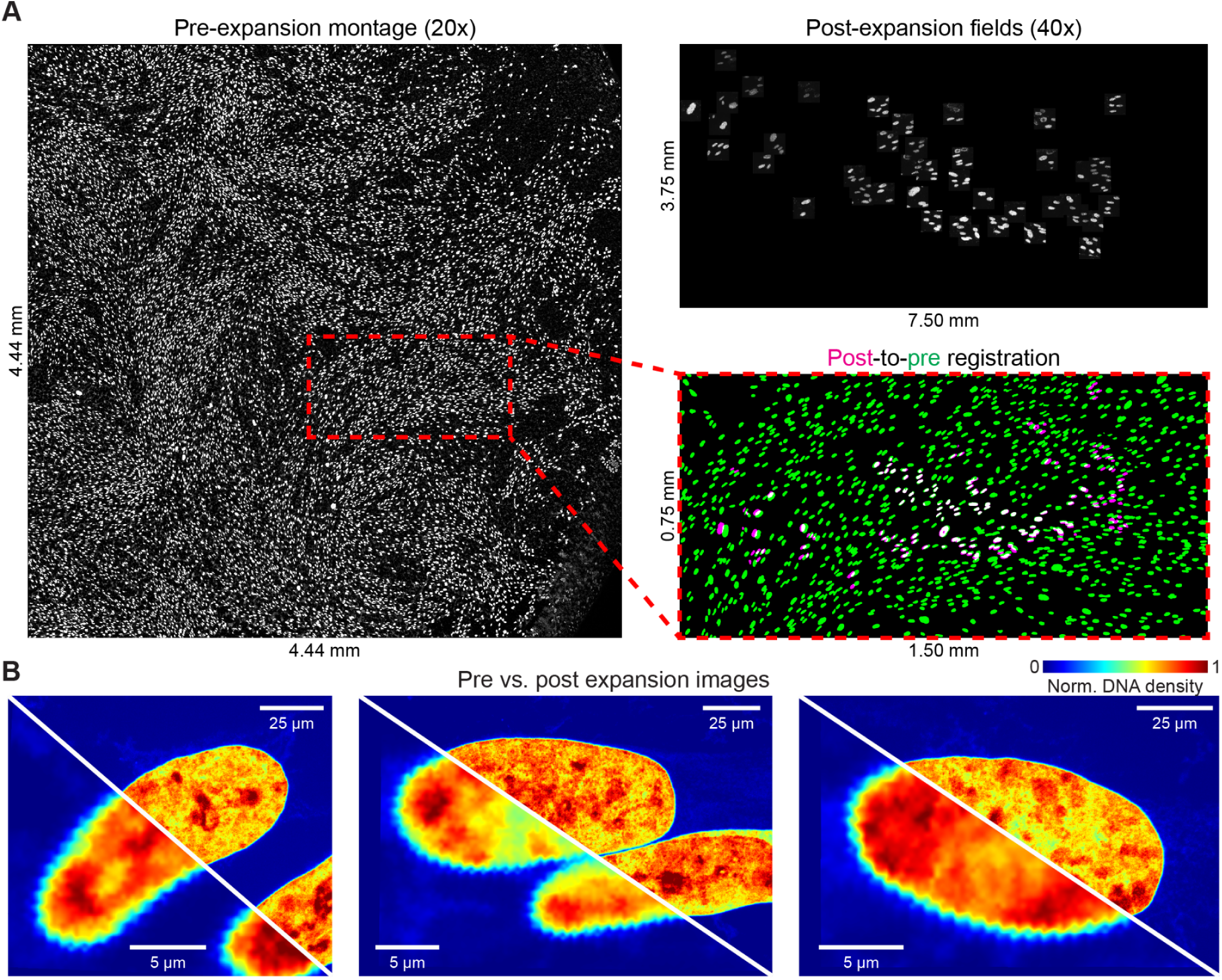
Quantification of expansion factor. (A) Left, DNA density (SYTOX Green) image montage of IMR-90 cells on a coverslip taken at 20x resolution, prior to expansion. Top right, DNA density image montage of a subset of the same IMR-90 cells in a gel following expansion, with individual fields of view taken at 40x resolution and combined using field *xy* coordinates. Bottom right, registration of pre- and post-expansion image segmentations. Green represents cells imaged only prior to expansion, white represents cells imaged both before and after, magenta represents cells with imperfect initial registration, which are later corrected during nuclei registration (**B**). Observed image sizes as marked. (B) Nuclei registration of pre-(bottom left half) and post-expansion (top right half) DNA density images. Scale bars represent observed distances. Sample shown had an expansion factor of approximately 5.

**Fig. S7:**
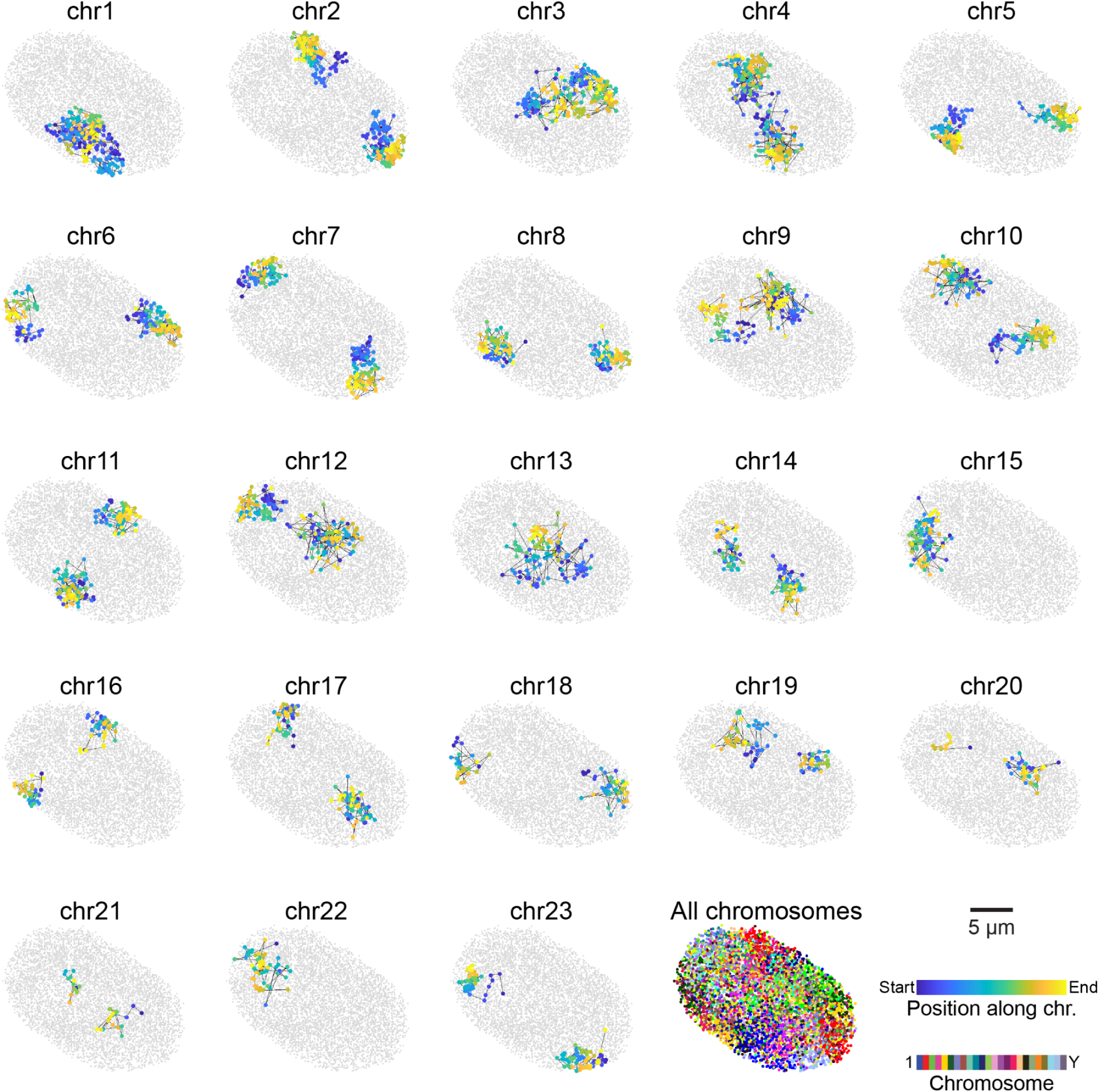
Visualization of chromosome territories in a skin fibroblast. ExIGS genomic reads for each chromosome in a skin fibroblast. Reads from each chromosome were assigned to one of two homologs for all autosomes and chrX (the donor of this cell line was female) based on spatial and genomic position. Colored dots indicate the position of the ExIGS reads on the given chromosome, gray dots represent ExIGS reads from other chromosomes. Black lines trace the 3D path each homolog takes through the nucleus. Shown as a maximum intensity projection in *z* for visualization. Scale bar in bottom right applies to all images, 5 microns.

**Fig. S8:**
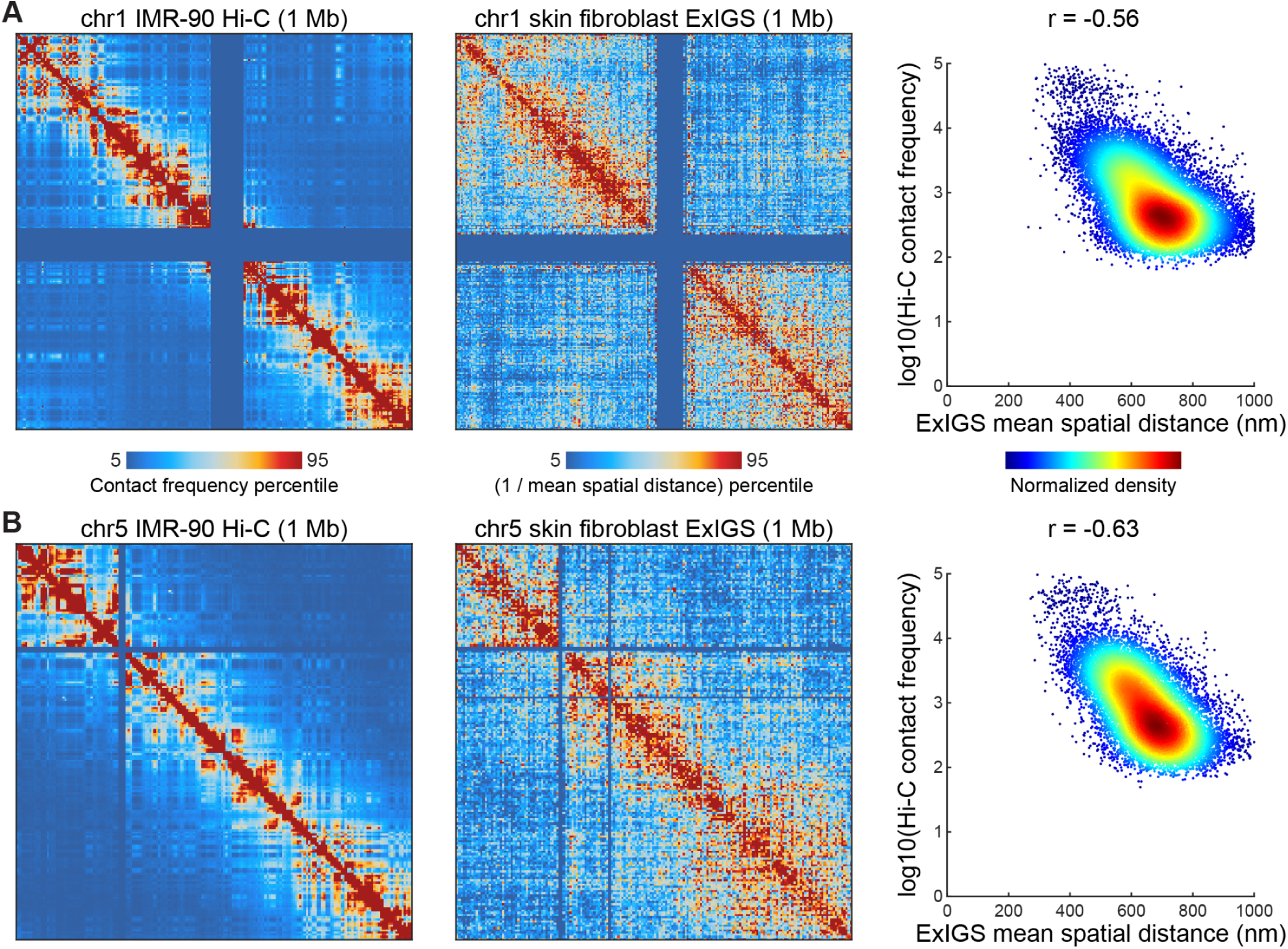
Comparison to Hi-C data. (A) Left, Contact frequency matrix for chr1 from Hi-C of IMR-90 fibroblasts from GSE63525(*92*) at 1 Mb resolution with Knight-Ruiz matrix balancing normalization. Middle, Inverse mean spatial distance matrix for chr1 from ExIGS of skin fibroblasts at 1 Mb resolution. Right, comparison of ExIGS mean spatial distance and normalized Hi-C contact frequency. Each point represents a pair of 1 Mb genomic bins from the chromosome shown. (B) Same as (A), but for chr5.

**Fig. S9:**
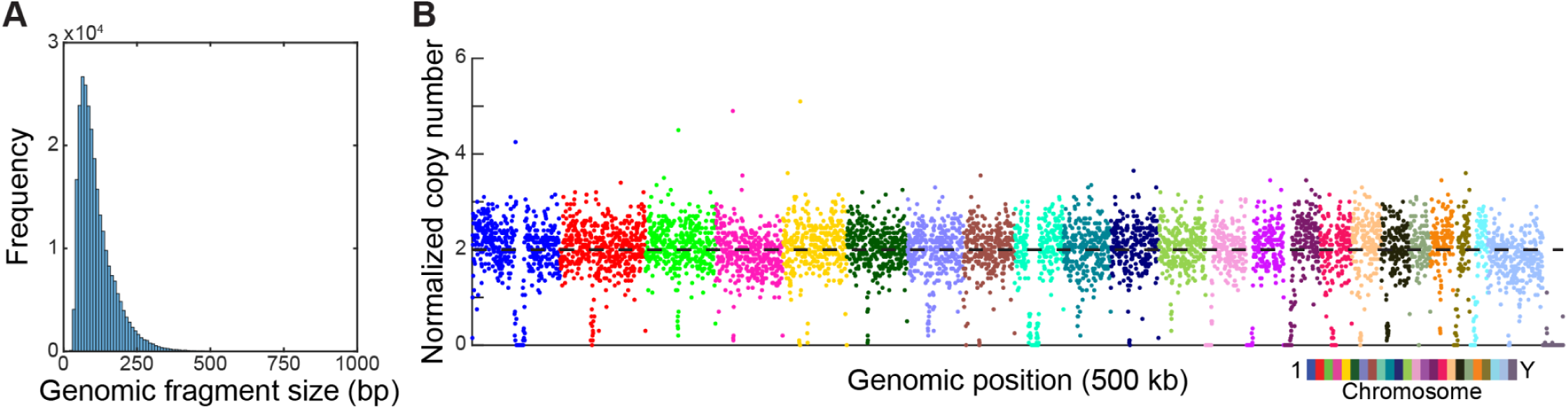
DNA fragment resolution and genomic coverage. (A) A histogram showing the genomic fragment sizes of all uniquely aligned ExIGS reads from skin fibroblasts. The peak of the distribution (∼100 bp) represents the average per-read genomic resolution of the method. (B) Normalized genomic coverage of non-overlapping 500 kb genomic bins of ExIGS reads from skin fibroblasts. Copy number was calculated by dividing by the median coverage and multiplying by 2. 96.4% of autosomal bins with more than 10 reads displayed a copy number between 1 and 3. Moderate peaks and valleys likely represent GC bias, points around zero likely fall in low mappability regions. Colors represent bin chromosome.

**Fig. S10:**
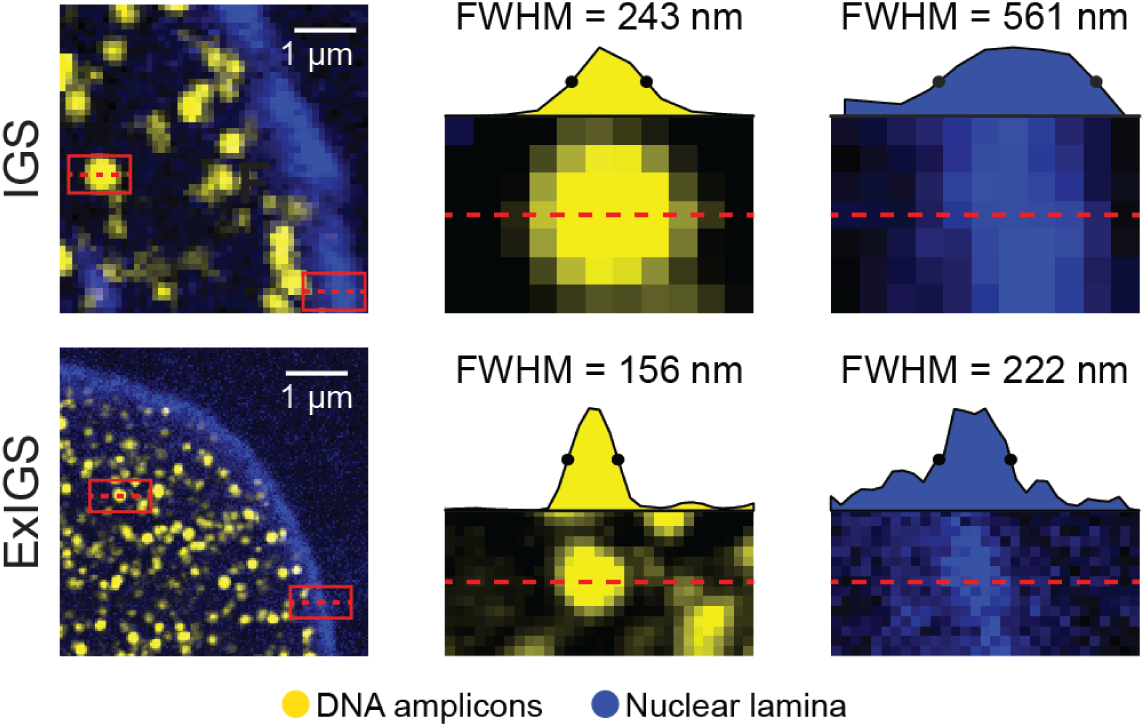
Quantification of expanded resolution. Top, Quantification of 3D amplicon and nuclear lamina feature width (lamin B) from *in situ* genome sequencing (IGS) of early mouse embryos. Bottom, Same as top, but for ExIGS of human skin fibroblasts (Lamin A/C). Feature widths were calculated using full width at half maximum measurements. The improvement in resolution is likely greater for lamin than DNA amplicons because the latter is not a diffraction-limited measurement (because amplicons are randomly imaged in 1 of 4 sequencing channels), but may also be due to species (mouse vs. human) or lamin type (Lamin B vs. Lamin A/C) differences.

**Fig. S11:**
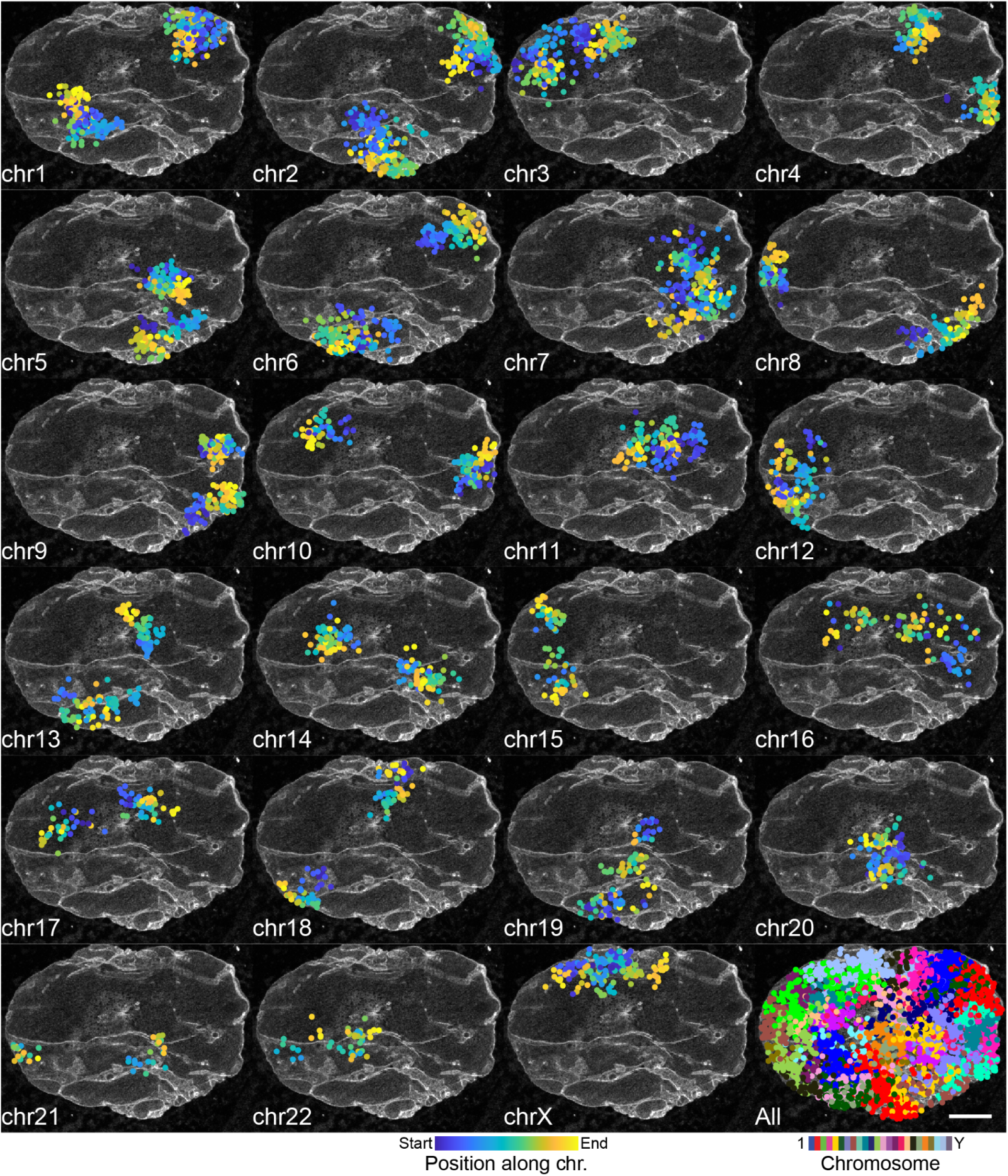
Visualization of chromosome territories and lamin abnormalities in a progeria fibroblast. ExIGS genomic reads for each chromosome in a passage 25 progeria fibroblast, overlaid on expansion immunofluorescence images of Lamin A/C. Both ExIGS reads and IF images are shown as maximum intensity *z*-projections for visualization. Scale bar in bottom right applies to all images, 5 microns.

**Fig. S12:**
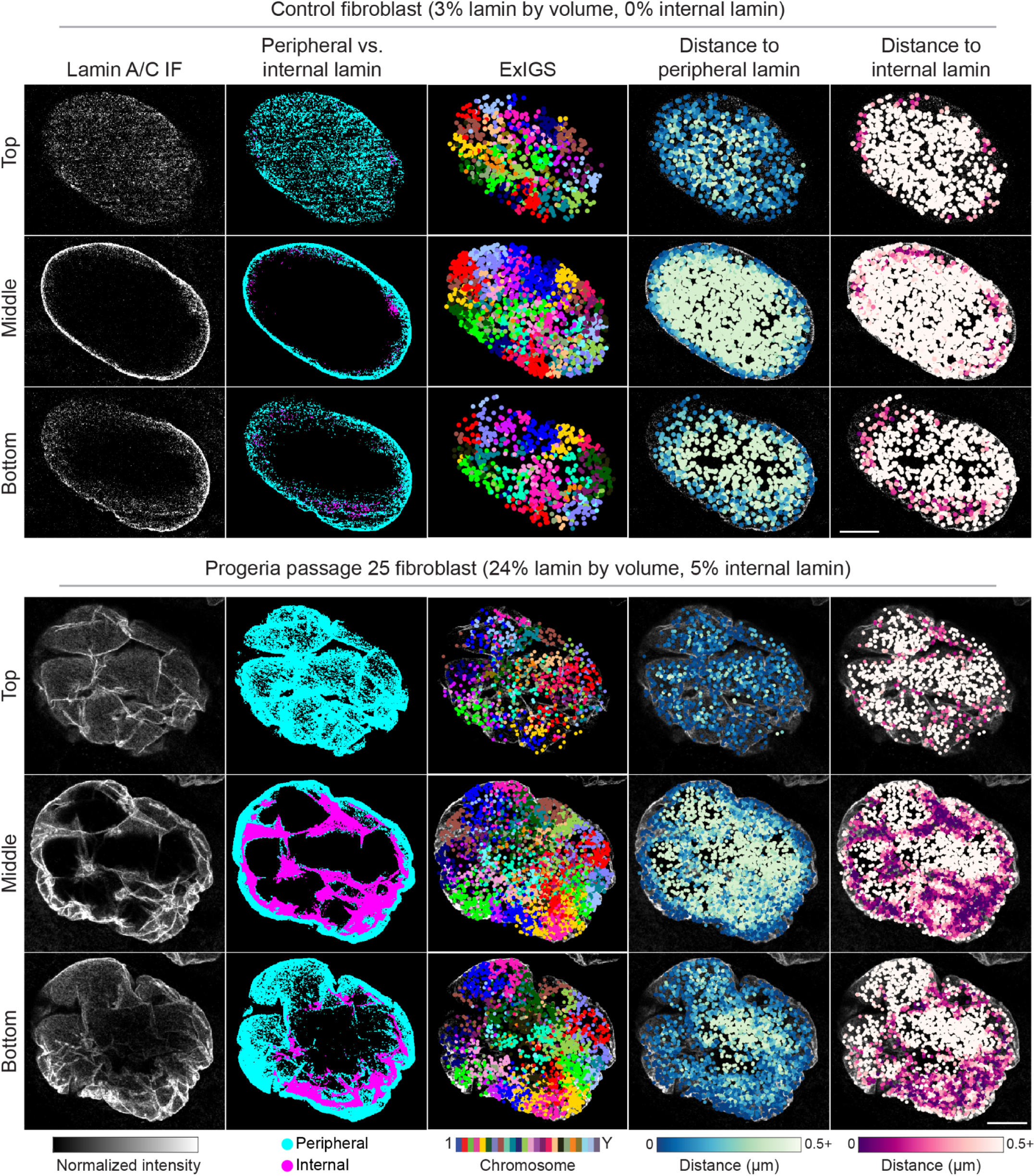
Segmentation of peripheral and internal lamin. Top, Lamin A/C segmentation and annotation of ExIGS reads by distance to lamin at top, middle, and bottom thirds of a control fibroblast. Bottom, Same as top, but for a passage 25 progeria fibroblast with substantial internal lamin. Both ExIGS reads and IF images are shown as maximum intensity *z*-projections of each third for visualization. Scale bar in bottom right applies to all images, 5 microns.

**Fig. S13:**
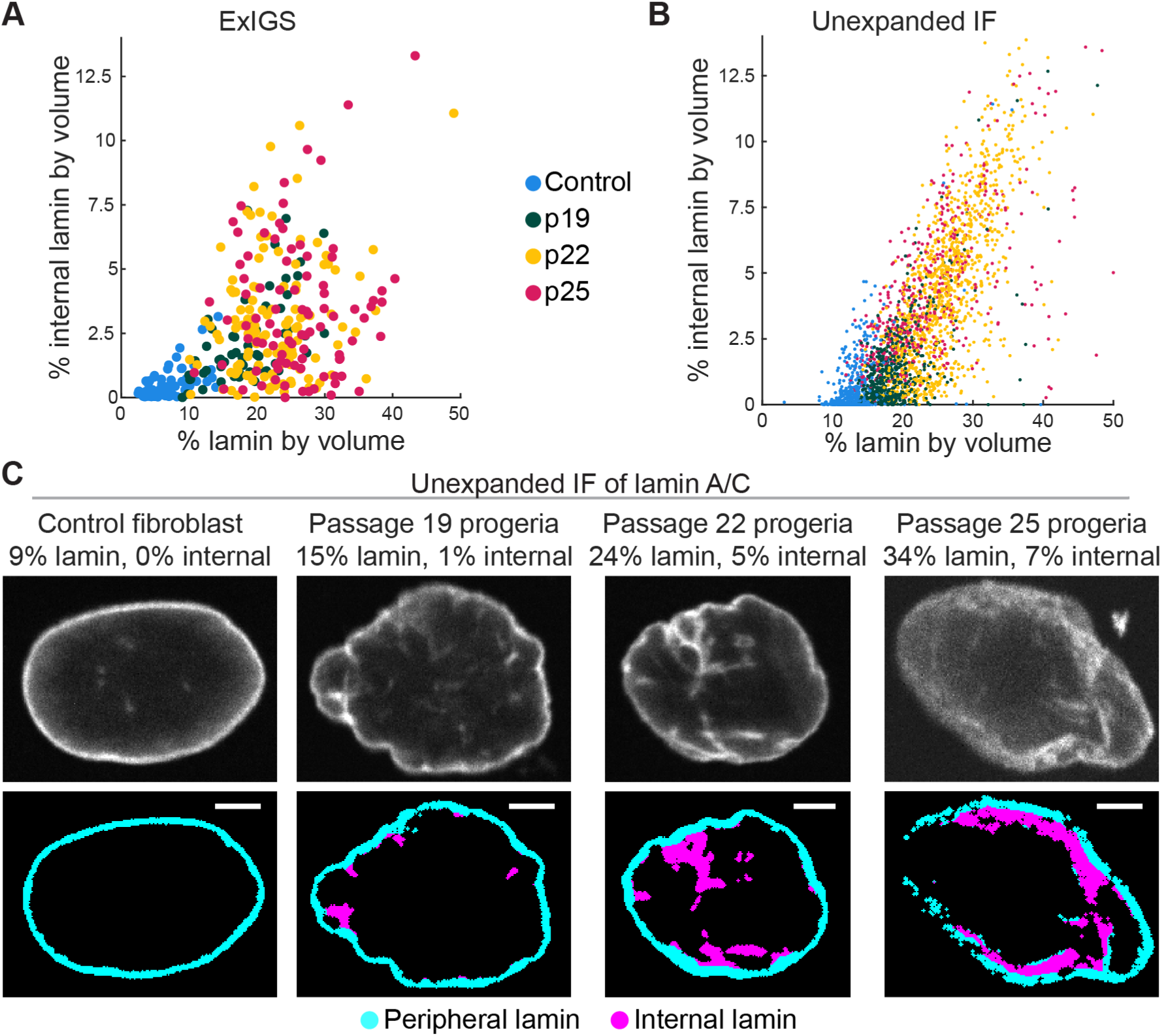
Validation of Lamin A/C accumulation in unexpanded progeria fibroblasts. (A) Reproduced from Fig. 2G. Control and progeria fibroblasts profiled with ExIGS and expansion immunofluorescence (IF) imaging of Lamin A/C, plotted by percent lamin by volume and percent internal lamin by volume. (B) Same as (A), but for non-expansion IF control and progeria fibroblasts from matched passages. (C) Top, Non-expansion IF of Lamin A/C for representative control and progeria fibroblasts from each passage. Bottom, segmentation of peripheral and internal lamin for the cells shown above. Scale bars, 5 microns. All images shown as single *z*-planes for visualization.

**Fig. S14:**
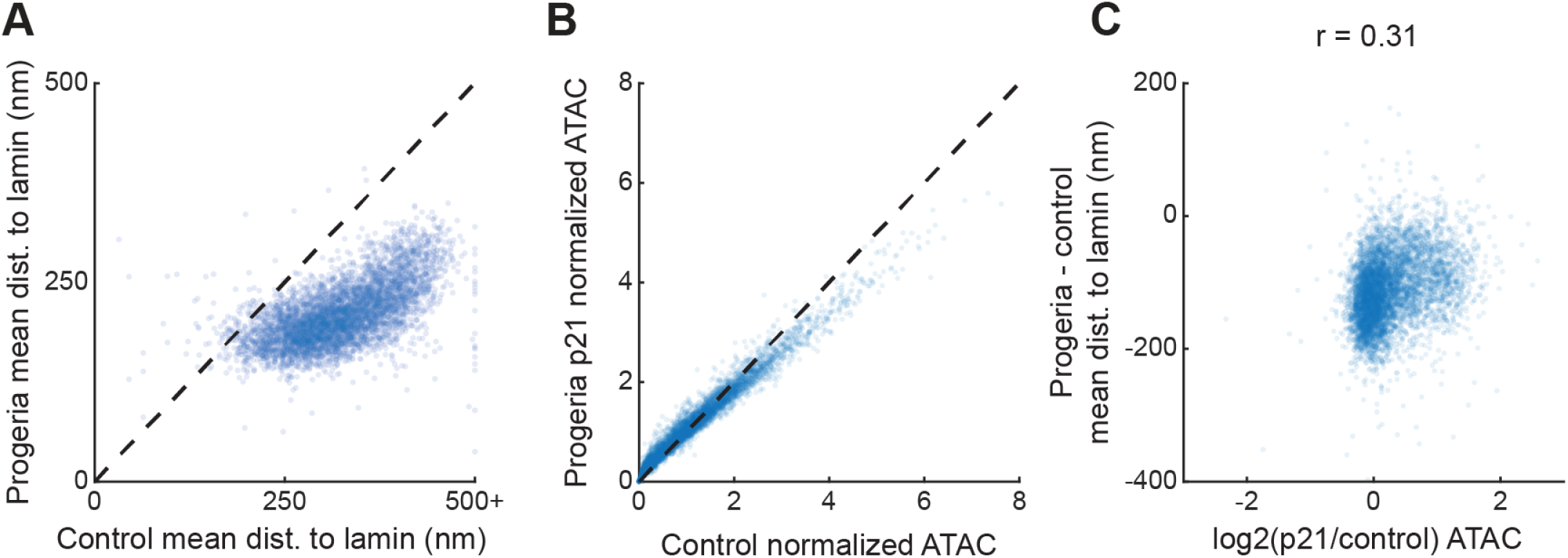
Concordant epigenetic changes in progeria fibroblasts. (A) Scatter plot showing mean distance of ExIGS reads to lamin in control fibroblasts vs. progeria (all passages). Each dot represents a non-overlapping 500 kb genomic bin. Dotted line indicates bins where mean distance to lamin is roughly the same. (B) Scatter plot showing normalized bulk ATAC-seq in control fibroblasts vs. passage 21 progeria fibroblasts. As in (A), each dot represents a 500 kb genomic bin and dotted line indicates similarity across conditions. ATAC-seq values are normalized by the mean over all bins in each condition. (C) Scatter plot showing the relationship between changes in normalized ATAC-seq and mean distance to lamin from control to progeria fibroblasts. The x-axis displays log2 fold changes using values from (B), while the y-axis displays differences usings values from (A). Positive correlation indicates that decreases in chromatin accessibility (negative values on the x-axis) are associated with closer proximity to the lamin (negative values on the y-axis).

**Fig. S15:**
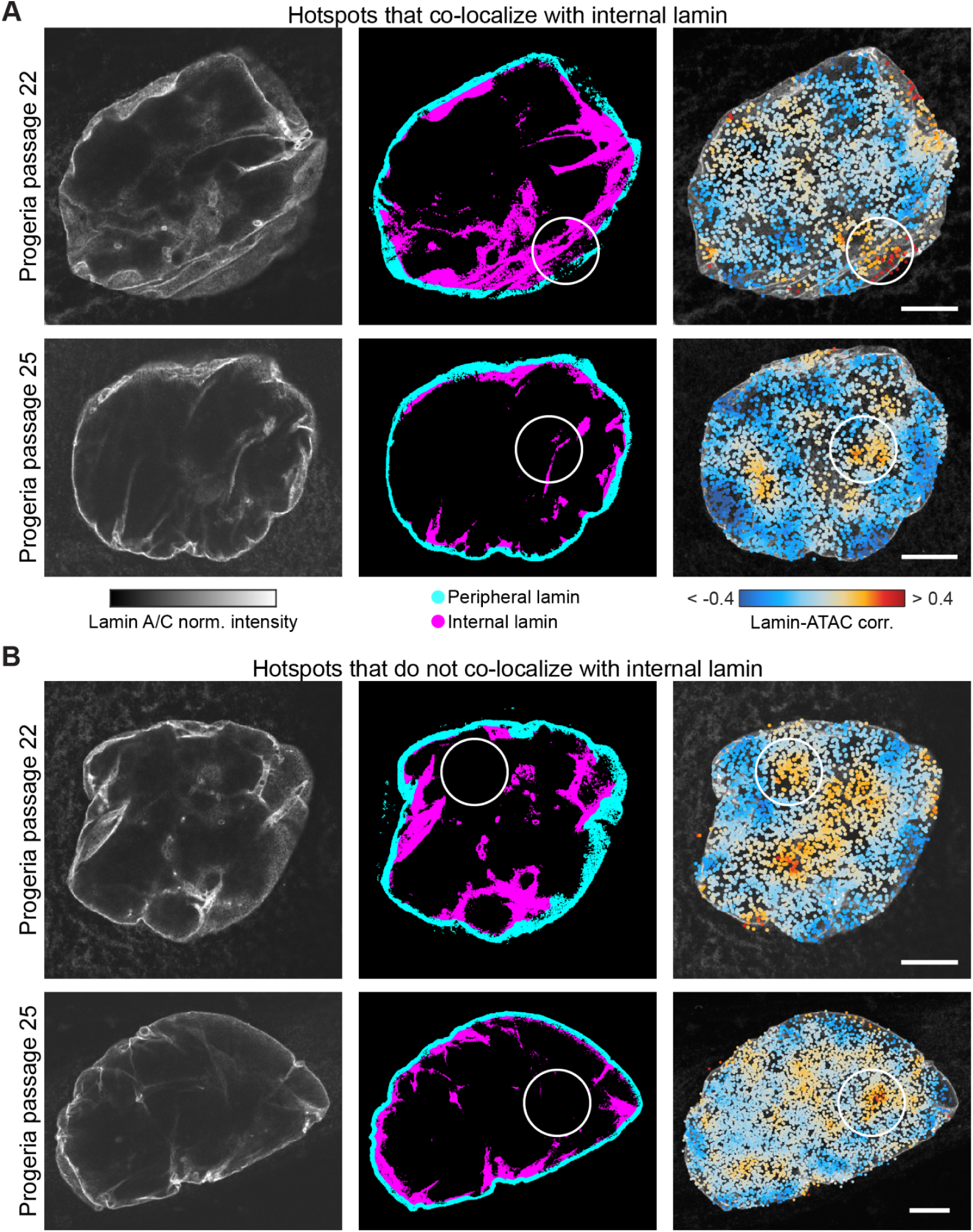
Stochastic co-localization of disruption hotspots and internal lamin. (A) Representative progeria fibroblasts from passage 22 (top) and 25 (bottom) with disruption hotspots that closely co-localize with internal lamin marked by white circles. Left and middle columns show a single z-plane for visualization, containing the lamin IF and segmentation. The right column shows ExIGS reads in a maximum intensity *z*-projection for a 1.8 micron thick section, labeled by hotspots. (B) Same as (A), but with disruption hotspots that do not co-localize with internal lamin are marked. All scale bars, 5 microns.

**Fig. S16:**
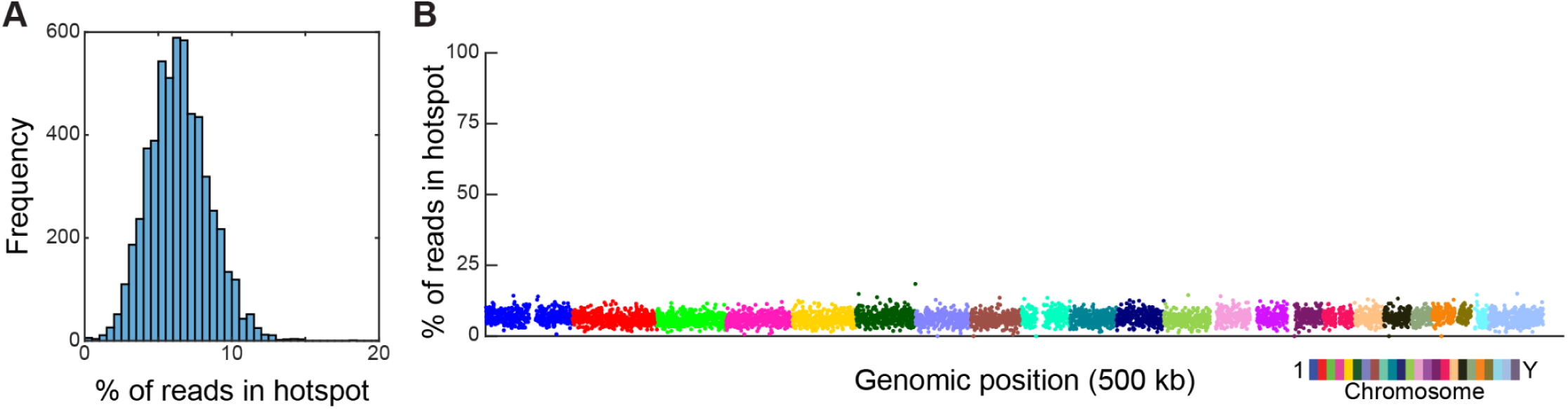
Disruption hotspots show random genomic distribution. (A) Histogram of percentage of reads falling within a disruption hotspot for non-overlapping 500 kb genomic bins. (B) Percentage of reads falling within a disruption hotspot by genomic position. Each dot represents a non-overlapping 500 kb genomic bin.

**Fig. S17:**
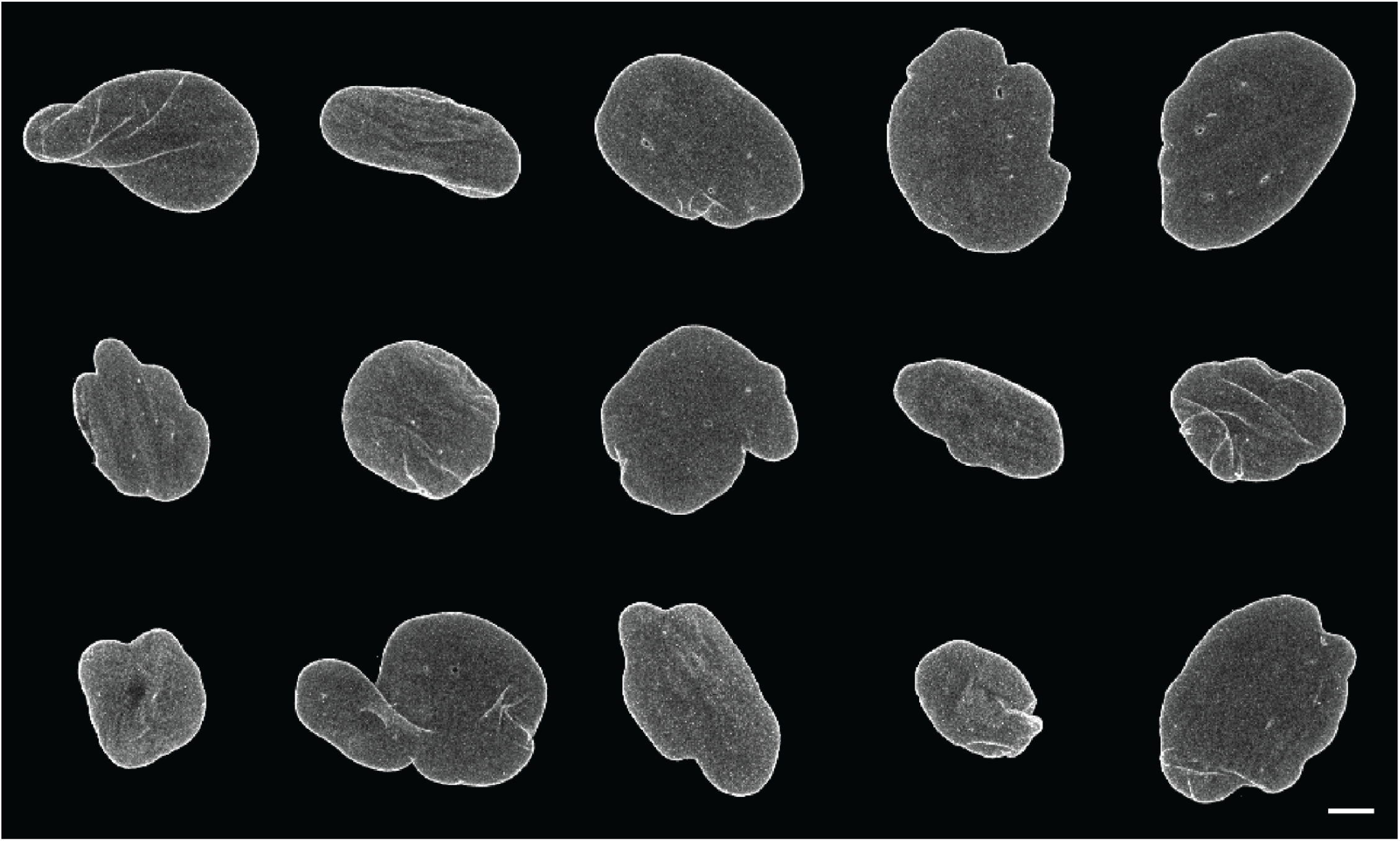
Visualization of nuclear abnormalities in fibroblasts from a 92 year-old donor. Expansion immunofluorescence (IF) images of Lamin A/C for 92 year-old donor fibroblasts with nuclear abnormalities. All images are 3D stacks, but are shown as maximum intensity z-projections for visualization purposes. Scale bar in bottom right applies to all nuclei, 5 microns.

## References

1. D. Zink, A. H. Fischer, J. A. Nickerson, Nuclear structure in cancer cells. Nat. Rev. Cancer 4 (2004).

2. M. Zwerger, C. Y. Ho, J. Lammerding, Nuclear Mechanics in Disease. Annu. Rev. Biomed. Eng. 13, 397 (2011).

3. J. R. Moffitt, E. Lundberg, H. Heyn, The emerging landscape of spatial profiling technologies. Nat. Rev. Genet. 23, 741–759 (2022).

4. A. Rao, D. Barkley, G. S. França, I. Yanai, Exploring tissue architecture using spatial transcriptomics. Nature 596, 211–220 (2021).

5. I. Jerkovic, G. Cavalli, Understanding 3D genome organization by multidisciplinary methods. Nat. Rev. Mol. Cell Biol. 22, 511–528 (2021).

6. A. Hafner, A. Boettiger, The spatial organization of transcriptional control. Nat. Rev. Genet. 24, 53–68 (2023).

7. A. Janssen, S. U. Colmenares, G. H. Karpen, Heterochromatin: Guardian of the Genome. Annu. Rev. Cell Dev. Biol. 34 (2018).

8. L. Ramos-Alonso, P. Holland, S. Le Gras, X. Zhao, B. Jost, M. Bjørås, Y. Barral, J. M. Enserink, P. Chymkowitch, Mitotic chromosome condensation resets chromatin to safeguard transcriptional homeostasis during interphase. Proc. Natl. Acad. Sci. U. S. A. 120 (2023).

9. Mechanosensing by the Lamina Protects against Nuclear Rupture, DNA Damage, and Cell-Cycle Arrest. Dev. Cell 49, 920–935.e5 (2019).

10. S. E. Johnstone, A. Reyes, Y. Qi, C. Adriaens, E. Hegazi, K. Pelka, J. H. Chen, L. S. Zou, Y. Drier, V. Hecht, N. Shoresh, M. K. Selig, C. A. Lareau, S. Iyer, S. C. Nguyen, E. F. Joyce, N. Hacohen, R. A. Irizarry, B. Zhang, M. J. Aryee, B. E. Bernstein, Large-Scale Topological Changes Restrain Malignant Progression in Colorectal Cancer. Cell 182, 1474–1489.e23 (2020).

11. L. Tan, J. Shi, S. Moghadami, B. Parasar, C. P. Wright, Y. Seo, K. Vallejo, I. Cobos, L. Duncan, R. Chen, K. Deisseroth, Lifelong restructuring of 3D genome architecture in cerebellar granule cells. Science 381, 1112–1119 (2023).

12. M. A. Ricci, C. Manzo, M. F. García-Parajo, M. Lakadamyali, M. P. Cosma, Chromatin fibers are formed by heterogeneous groups of nucleosomes in vivo. Cell 160, 1145–1158 (2015).

13. A. N. Boettiger, B. Bintu, J. R. Moffitt, S. Wang, B. J. Beliveau, G. Fudenberg, M. Imakaev, L. A. Mirny, C.-T. Wu, X. Zhuang, Super-resolution imaging reveals distinct chromatin folding for different epigenetic states. Nature 529, 418–422 (2016).

14. H. D. Ou, S. Phan, T. J. Deerinck, A. Thor, M. H. Ellisman, C. C. O’Shea, ChromEMT: Visualizing 3D chromatin structure and compaction in interphase and mitotic cells. Science 357 (2017).

15. T. Nozaki, R. Imai, M. Tanbo, R. Nagashima, S. Tamura, T. Tani, Y. Joti, M. Tomita, K. Hibino, M. T. Kanemaki, K. S. Wendt, Y. Okada, T. Nagai, K. Maeshima, Dynamic Organization of Chromatin Domains Revealed by Super-Resolution Live-Cell Imaging. Mol. Cell 67, 282–293.e7 (2017).

16. E. Miron, R. Oldenkamp, J. M. Brown, D. M. S. Pinto, C. S. Xu, A. R. Faria, H. A. Shaban, J. D. P. Rhodes, C. Innocent, S. de Ornellas, H. F. Hess, V. Buckle, L. Schermelleh, Chromatin arranges in chains of mesoscale domains with nanoscale functional topography independent of cohesin. Sci Adv 6 (2020).

17. L. Xie, P. Dong, X. Chen, T.-H. S. Hsieh, S. Banala, M. De Marzio, B. P. English, Y. Qi, S. K. Jung, K.-R. Kieffer-Kwon, W. R. Legant, A. S. Hansen, A. Schulmann, R. Casellas, B. Zhang, E. Betzig, L. D. Lavis, H. Y. Chang, R. Tjian, Z. Liu, 3D ATAC-PALM: super-resolution imaging of the accessible genome. Nat. Methods 17, 430–436 (2020).

18. M. Gelléri, S.-Y. Chen, B. Hübner, J. Neumann, O. Kröger, F. Sadlo, J. Imhoff, M. J. Hendzel, M. Cremer, T. Cremer, H. Strickfaden, C. Cremer, True-to-scale DNA-density maps correlate with major accessibility differences between active and inactive chromatin. Cell Rep. 42, 112567 (2023).

19. J. Xu, H. Ma, J. Jin, S. Uttam, R. Fu, Y. Huang, Y. Liu, Super-Resolution Imaging of Higher-Order Chromatin Structures at Different Epigenomic States in Single Mammalian Cells. Cell Rep. 24, 873–882 (2018).

20. H. Q. Nguyen, S. Chattoraj, D. Castillo, S. C. Nguyen, G. Nir, A. Lioutas, E. A. Hershberg, N. M. C. Martins, P. L. Reginato, M. Hannan, B. J. Beliveau, G. M. Church, E. R. Daugharthy, M. A. Marti-Renom, C.-T. Wu, 3D mapping and accelerated super-resolution imaging of the human genome using in situ sequencing. Nat. Methods 17, 822–832 (2020).

21. J.-H. Su, P. Zheng, S. S. Kinrot, B. Bintu, X. Zhuang, Genome-Scale Imaging of the 3D Organization and Transcriptional Activity of Chromatin. Cell 182, 1641–1659.e26 (2020).

22. Y. Takei, J. Yun, S. Zheng, N. Ollikainen, N. Pierson, J. White, S. Shah, J. Thomassie, S. Suo, C.-H. L. Eng, M. Guttman, G.-C. Yuan, L. Cai, Integrated spatial genomics reveals global architecture of single nuclei. Nature 590, 344–350 (2021).

23. Y. Takei, S. Zheng, J. Yun, S. Shah, N. Pierson, J. White, S. Schindler, C. H. Tischbirek, G.-C. Yuan, L. Cai, Single-cell nuclear architecture across cell types in the mouse brain. Science 374, 586–594 (2021).

24. T. Lu, C. E. Ang, X. Zhuang, Spatially resolved epigenomic profiling of single cells in complex tissues. Cell 185, 4448–4464.e17 (2022).

25. A. C. Payne, Z. D. Chiang, P. L. Reginato, S. M. Mangiameli, E. M. Murray, C.-C. Yao, S. Markoulaki, A. S. Earl, A. S. Labade, R. Jaenisch, G. M. Church, E. S. Boyden, J. D. Buenrostro, F. Chen, In situ genome sequencing resolves DNA sequence and structure in intact biological samples. Science 371 (2021).

26. F. Chen, P. W. Tillberg, E. S. Boyden, Optical imaging. Expansion microscopy. Science 347, 543–548 (2015).

27. M. A. Woodworth, K. K. H. Ng, A. R. Halpern, N. A. Pease, P. H. B. Nguyen, H. Y. Kueh, J. C. Vaughan, Multiplexed single-cell profiling of chromatin states at genomic loci by expansion microscopy. Nucleic Acids Res. 49, e82 (2021).

28. M. E. Pownall, L. Miao, C. E. Vejnar, O. M’Saad, A. Sherrard, M. A. Frederick, M. D. J. Benitez, C. W. Boswell, K. S. Zaret, J. Bewersdorf, A. J. Giraldez, Chromatin expansion microscopy reveals nanoscale organization of transcription and chromatin. Science 381, 92–100 (2023).

29. F. Chen, A. T. Wassie, A. J. Cote, A. Sinha, S. Alon, S. Asano, E. R. Daugharthy, J.-B. Chang, A. Marblestone, G. M. Church, A. Raj, E. S. Boyden, Nanoscale imaging of RNA with expansion microscopy. Nat. Methods 13, 679–684 (2016).

30. S. Alon, D. R. Goodwin, A. Sinha, A. T. Wassie, F. Chen, E. R. Daugharthy, Y. Bando, A. Kajita, A. G. Xue, K. Marrett, R. Prior, Y. Cui, A. C. Payne, C.-C. Yao, H.-J. Suk, R. Wang, C.-C. J. Yu, P. Tillberg, P. Reginato, N. Pak, S. Liu, S. Punthambaker, E. P. R. Iyer, R. E. Kohman, J. A. Miller, E. S. Lein, A. Lako, N. Cullen, S. Rodig, K. Helvie, D. L. Abravanel, N. Wagle, B. E. Johnson, J. Klughammer, M. Slyper, J. Waldman, J. Jané-Valbuena, O. Rozenblatt-Rosen, A. Regev, IMAXT Consortium, G. M. Church, A. H. Marblestone, E. S. Boyden, Expansion sequencing: Spatially precise in situ transcriptomics in intact biological systems. Science 371 (2021).

31. A. Klimas, B. R. Gallagher, P. Wijesekara, S. Fekir, E. F. DiBernardo, Z. Cheng, D. B. Stolz, F. Cambi, S. C. Watkins, S. L. Brody, A. Horani, A. L. Barth, C. I. Moore, X. Ren, Y. Zhao, Magnify is a universal molecular anchoring strategy for expansion microscopy. Nat. Biotechnol. 41, 858–869 (2023).

32. H. Li, A. R. Warden, J. He, G. Shen, X. Ding, Expansion microscopy with ninefold swelling (NIFS) hydrogel permits cellular ultrastructure imaging on conventional microscope. Science Advances, doi: 10.1126/sciadv.abm4006 (2022).

33. X. Chen, Y. Shen, W. Draper, J. D. Buenrostro, U. Litzenburger, S. W. Cho, A. T. Satpathy, A. C. Carter, R. P. Ghosh, A. East-Seletsky, J. A. Doudna, W. J. Greenleaf, J. T. Liphardt, H. Y. Chang, ATAC-see reveals the accessible genome by transposase-mediated imaging and sequencing. Nat. Methods 13, 1013–1020 (2016).

34. D. Feldman, A. Singh, J. L. Schmid-Burgk, R. J. Carlson, A. Mezger, A. J. Garrity, F. Zhang, P. C. Blainey, Optical Pooled Screens in Human Cells. Cell 179 (2019).

35. T. Cremer, M. Cremer, Chromosome territories. *Cold Sprin*g Harb. Perspect. Biol. 2 (2010).

36. Roadmap Epigenomics Consortium, A. Kundaje, W. Meuleman, J. Ernst, M. Bilenky, A. Yen, A. Heravi-Moussavi, P. Kheradpour, Z. Zhang, J. Wang, M. J. Ziller, V. Amin, J. W. Whitaker, M. D. Schultz, L. D. Ward, A. Sarkar, G. Quon, R. S. Sandstrom, M. L. Eaton, Y.-C. Wu, A. R. Pfenning, X. Wang, M. Claussnitzer, Y. Liu, C. Coarfa, R. A. Harris, N. Shoresh, C. B. Epstein, E. Gjoneska, D. Leung, W. Xie, R. D. Hawkins, R. Lister, C. Hong, P. Gascard, A. J. Mungall, R. Moore, E. Chuah, A. Tam, T. K. Canfield, R. S. Hansen, R. Kaul, P. J. Sabo, M. S. Bansal, A. Carles, J. R. Dixon, K.-H. Farh, S. Feizi, R. Karlic, A.-R. Kim, A. Kulkarni, D. Li, R. Lowdon, G. Elliott, T. R. Mercer, S. J. Neph, V. Onuchic, P. Polak, N. Rajagopal, P. Ray, R. C. Sallari, K. T. Siebenthall, N. A. Sinnott-Armstrong, M. Stevens, R. E. Thurman, J. Wu, B. Zhang, X. Zhou, A. E. Beaudet, L. A. Boyer, P. L. De Jager, P. J. Farnham, S. J. Fisher, D. Haussler, S. J. M. Jones, W. Li, M. A. Marra, M. T. McManus, S. Sunyaev, J. A. Thomson, T. D. Tlsty, L.-H. Tsai, W. Wang, R. A. Waterland, M. Q. Zhang, L. H. Chadwick, B. E. Bernstein, J. F. Costello, J. R. Ecker, M. Hirst, A. Meissner, A. Milosavljevic, B. Ren, J. A. Stamatoyannopoulos, T. Wang, M. Kellis, Integrative analysis of 111 reference human epigenomes. Nature 518, 317–330 (2015).

37. B. Huang, H. Babcock, X. Zhuang, Breaking the diffraction barrier: super-resolution imaging of cells. Cell 143, 1047–1058 (2010).

38. R. de Leeuw, Y. Gruenbaum, O. Medalia, Nuclear Lamins: Thin Filaments with Major Functions. Trends Cell Biol. 28, 34–45 (2018).

39. B. Nmezi, J. Xu, R. Fu, T. J. Armiger, G. Rodriguez-Bey, J. S. Powell, H. Ma, M. Sullivan, Y. Tu, N. Y. Chen, S. G. Young, D. B. Stolz, K. N. Dahl, Y. Liu, Q. S. Padiath, Concentric organization of A- and B-type lamins predicts their distinct roles in the spatial organization and stability of the nuclear lamina. Proc. Natl. Acad. Sci. U. S. A. 116, 4307–4315 (2019).

40. B. van Steensel, A. S. Belmont, Lamina-associated domains: links with chromosome architecture, heterochromatin and gene repression. Cell 169, 780 (2017).

41. N. Briand, P. Collas, Lamina-associated domains: peripheral matters and internal affairs. Genome Biol. 21, 85 (2020).

42. K. L. Reddy, J. M. Zullo, E. Bertolino, H. Singh, Transcriptional repression mediated by repositioning of genes to the nuclear lamina. Nature 452, 243–247 (2008).

43. H. Wang, X. Xu, C. M. Nguyen, Y. Liu, Y. Gao, X. Lin, T. Daley, N. H. Kipniss, M. La Russa, L. S. Qi, CRISPR-Mediated Programmable 3D Genome Positioning and Nuclear Organization. Cell 175, 1405–1417.e14 (2018).

44. J. M. Bridger, I. R. Kill, M. O’Farrell, C. J. Hutchison, Internal lamin structures within G1 nuclei of human dermal fibroblasts. J. Cell Sci. 104 (Pt 2), 297–306 (1993).

45. N. Naetar, S. Ferraioli, R. Foisner, Lamins in the nuclear interior - life outside the lamina. J. Cell Sci. 130 (2017).

46. K. Gesson, P. Rescheneder, M. P. Skoruppa, A. von Haeseler, T. Dechat, R. Foisner, A-type lamins bind both hetero- and euchromatin, the latter being regulated by lamina-associated polypeptide 2 alpha. Genome Res. 26, 462–473 (2016).

47. K. Ikegami, S. Secchia, O. Almakki, J. D. Lieb, I. P. Moskowitz, Phosphorylated Lamin A/C in the Nuclear Interior Binds Active Enhancers Associated with Abnormal Transcription in Progeria. Dev. Cell 52, 699–713.e11 (2020).

48. P. P. Shah, K. C. Keough, K. Gjoni, G. T. Santini, R. J. Abdill, N. M. Wickramasinghe, C. E. Dundes, A. Karnay, A. Chen, R. E. A. Salomon, P. J. Walsh, S. C. Nguyen, S. Whalen, E. F. Joyce, K. M. Loh, N. Dubois, K. S. Pollard, R. Jain, An atlas of lamina-associated chromatin across twelve human cell types reveals an intermediate chromatin subtype. Genome Biol. 24, 16 (2023).

49. J. Kind, B. van Steensel, Stochastic genome-nuclear lamina interactions: modulating roles of Lamin A and BAF. Nucleus 5, 124–130 (2014).

50. B. C. Capell, F. S. Collins, Human laminopathies: nuclei gone genetically awry. Nat. Rev. Genet. 7, 940–952 (2006).

51. L. B. Gordon, W. Ted Brown, F. S. Collins, Hutchinson-Gilford Progeria Syndrome (University of Washington, Seattle, 2023).

52. R. D. Goldman, D. K. Shumaker, M. R. Erdos, M. Eriksson, A. E. Goldman, L. B. Gordon, Y. Gruenbaum, S. Khuon, M. Mendez, R. Varga, F. S. Collins, Accumulation of mutant lamin A causes progressive changes in nuclear architecture in Hutchinson-Gilford progeria syndrome. Proc. Natl. Acad. Sci. U. S. A. 101, 8963–8968 (2004).

53. D. K. Shumaker, T. Dechat, A. Kohlmaier, S. A. Adam, M. R. Bozovsky, M. R. Erdos, M. Eriksson, A. E. Goldman, S. Khuon, F. S. Collins, T. Jenuwein, R. D. Goldman, Mutant nuclear lamin A leads to progressive alterations of epigenetic control in premature aging. Proc. Natl. Acad. Sci. U. S. A. 103, 8703–8708 (2006).

54. R. P. McCord, A. Nazario-Toole, H. Zhang, P. S. Chines, Y. Zhan, M. R. Erdos, F. S. Collins, J. Dekker, K. Cao, Correlated alterations in genome organization, histone methylation, and DNA-lamin A/C interactions in Hutchinson-Gilford progeria syndrome. Genome Res. 23, 260–269 (2013).

55. J. Perovanovic, S. Dell’Orso, V. F. Gnochi, J. K. Jaiswal, V. Sartorelli, C. Vigouroux, K. Mamchaoui, V. Mouly, G. Bonne, E. P. Hoffman, Laminopathies disrupt epigenomic developmental programs and cell fate. Sci. Transl. Med. 8, 335ra58 (2016).

56. F. Köhler, F. Bormann, G. Raddatz, J. Gutekunst, S. Corless, T. Musch, A. S. Lonsdorf, S. Erhardt, F. Lyko, M. Rodríguez-Paredes, Epigenetic deregulation of lamina-associated domains in Hutchinson-Gilford progeria syndrome. Genome Med. 12, 46 (2020).

57. E. Sebestyén, F. Marullo, F. Lucini, C. Petrini, A. Bianchi, S. Valsoni, I. Olivieri, L. Antonelli, F. Gregoretti, G. Oliva, F. Ferrari, C. Lanzuolo, SAMMY-seq reveals early alteration of heterochromatin and deregulation of bivalent genes in Hutchinson-Gilford Progeria Syndrome. Nat. Commun. 11, 6274 (2020).

58. P. P. Shah, W. Lv, J. H. Rhoades, A. Poleshko, D. Abbey, M. A. Caporizzo, R. Linares-Saldana, J. G. Heffler, N. Sayed, D. Thomas, Q. Wang, L. J. Stanton, K. Bedi, M. P. Morley, T. P. Cappola, A. T. Owens, K. B. Margulies, D. B. Frank, J. C. Wu, D. J. Rader, W. Yang, B. L. Prosser, K. Musunuru, R. Jain, Pathogenic LMNA variants disrupt cardiac lamina-chromatin interactions and de-repress alternative fate genes. Cell Stem Cell 28, 938–954.e9 (2021).

59. M. Chiang, D. Michieletto, C. A. Brackley, N. Rattanavirotkul, H. Mohammed, D. Marenduzzo, T. Chandra, Polymer Modeling Predicts Chromosome Reorganization in Senescence. Cell Rep. 28 (2019).

60. C. Y. McLean, D. Bristor, M. Hiller, S. L. Clarke, B. T. Schaar, C. B. Lowe, A. M. Wenger, G. Bejerano, GREAT improves functional interpretation of cis-regulatory regions. Nat. Biotechnol. 28, 495 (2010).

61. W. Chang, Y. Wang, G. W. G. Luxton, C. Östlund, H. J. Worman, G. G. Gundersen, Imbalanced nucleocytoskeletal connections create common polarity defects in progeria and physiological aging. Proc. Natl. Acad. Sci. U. S. A. 116 (2019).

62. X. Mu, C. Tseng, W. S. Hambright, P. Matre, C. Y. Lin, P. Chanda, W. Chen, J. Gu, S. Ravuri, Y. Cui, L. Zhong, J. P. Cooke, L. J. Niedernhofer, P. D. Robbins, J. Huard, Cytoskeleton stiffness regulates cellular senescence and innate immune response in Hutchinson-Gilford Progeria Syndrome. Aging Cell 19 (2020).

63. D. McClintock, D. Ratner, M. Lokuge, D. M. Owens, L. B. Gordon, F. S. Collins, K. Djabali, The mutant form of lamin A that causes Hutchinson-Gilford progeria is a biomarker of cellular aging in human skin. PLoS One 2 (2007).

64. P. Scaffidi, T. Misteli, Lamin A-Dependent Nuclear Defects in Human Aging. Science, doi: 10.1126/science.1127168 (2006).

65. K. Cao, B. C. Capell, M. R. Erdos, K. Djabali, F. S. Collins, A lamin A protein isoform overexpressed in Hutchinson-Gilford progeria syndrome interferes with mitosis in progeria and normal cells. Proc. Natl. Acad. Sci. U. S. A. 104 (2007).

66. A. Prakash, L. B. Gordon, M. E. Kleinman, E. B. Gurary, J. Massaro, R. D’Agostino Sr, M. W. Kieran, M. Gerhard-Herman, L. Smoot, Cardiac Abnormalities in Patients With Hutchinson-Gilford Progeria Syndrome. JAMA Cardiology 3, 326 (2018).

67. C. P. Martinez-Jimenez, N. Eling, H.-C. Chen, C. A. Vallejos, A. A. Kolodziejczyk, F. Connor, L. Stojic, T. F. Rayner, M. J. T. Stubbington, S. A. Teichmann, M. de la Roche, J. C. Marioni, D. T. Odom, Aging increases cell-to-cell transcriptional variability upon immune stimulation. Science 355, 1433 (2017).

68. R. Stegeman, V. M. Weake, Transcriptional signatures of aging. J. Mol. Biol. 429, 2427 (2017).

69. C. A. Schmitt, B. Wang, M. Demaria, Senescence and cancer - role and therapeutic opportunities. Nat. Rev. Clin. Oncol. 19, 619–636 (2022).

70. M. Shokrollahi, M. Stanic, A. Hundal, J. N. Y. Chan, D. Urman, C. A. Jordan, A. Hakem, R. Espin, J. Hao, R. Krishnan, P. G. Maass, B. C. Dickson, M. P. Hande, M. A. Pujana, R. Hakem, K. Mekhail, DNA double-strand break–capturing nuclear envelope tubules drive DNA repair. Nat. Struct. Mol. Biol., 1–12 (2024).

71. J.-H. Yang, M. Hayano, P. T. Griffin, J. A. Amorim, M. S. Bonkowski, E. L. Salfati, M. Blanchette, E. M. Munding, M. Bhakta, Y. C. Chew, W. Guo, X. Yang, S. Maybury-Lewis, X. Tian, J. M. Ross, G. Coppotelli, M. V. Meer, R. Rogers-Hammond, D. L. Vera, Y. R. Lu, J. W. Pippin, M. L. Creswell, Z. Dou, C. Xu, S. J. Mitchell, A. Das, B. L. O’Connell, S. Thakur, A. E. Kane, Q. Su, Y. Mohri, E. K. Nishimura, L. Schaevitz, N. Garg, A.-M. Balta, M. A. Rego, M. Gregory-Ksander, T. C. Jakobs, L. Zhong, H. Wakimoto, J. El Andari, D. Grimm, R. Mostoslavsky, A. J. Wagers, K. Tsubota, S. J. Bonasera, C. M. Palmeira, J. G. Seidman, C. E. Seidman, N. S. Wolf, J. A. Kreiling, J. M. Sedivy, G. F. Murphy, R. E. Green, B. A. Garcia, S. L. Berger, P. Oberdoerffer, S. J. Shankland, V. N. Gladyshev, B. R. Ksander, A. R. Pfenning, L. A. Rajman, D. A. Sinclair, Loss of epigenetic information as a cause of mammalian aging. Cell 186, 305–326.e27 (2023).

72. C. P. Martinez-Jimenez, N. Eling, H.-C. Chen, C. A. Vallejos, A. A. Kolodziejczyk, F. Connor, L. Stojic, T. F. Rayner, M. J. T. Stubbington, S. A. Teichmann, M. de la Roche, J. C. Marioni, D. T. Odom, Aging increases cell-to-cell transcriptional variability upon immune stimulation. Science 355, 1433 (2017).

73. C. Debès, A. Papadakis, S. Grönke, Ö. Karalay, L. S. Tain, A. Mizi, S. Nakamura, O. Hahn, C. Weigelt, N. Josipovic, A. Zirkel, I. Brusius, K. Sofiadis, M. Lamprousi, Y.-X. Lu, W. Huang, R. Esmaillie, T. Kubacki, M. R. Späth, B. Schermer, T. Benzing, R.-U. Müller, A. Antebi, L. Partridge, A. Papantonis, A. Beyer, Ageing-associated changes in transcriptional elongation influence longevity. Nature 616, 814–821 (2023).

74. A. B. C. Capell, M. R. Erdos, J. P. Madigan, J. J. Fiordalisi, R. Varga, K. N. Conneely, L. B. Gordon, C. J. Der, D. Cox, F. S. Collins, Inhibiting farnesylation of progerin prevents the characteristic nuclear blebbing of Hutchinson-Gilford progeria syndrome. Proc. Natl. Acad. Sci. U. S. A. 102 (2005).

74. A. Ocampo, P. Reddy, P. Martinez-Redondo, A. Platero-Luengo, F. Hatanaka, T. Hishida, M. Li, D. Lam, M. Kurita, E. Beyret, T. Araoka, E. Vazquez-Ferrer, D. Donoso, J. L. Roman, J. Xu, C. R. Esteban, G. Nuñez, E. N. Delicado, J. M. Campistol, I. Guillen, P. Guillen, J. C. I. Belmonte, In Vivo Amelioration of Age-Associated Hallmarks by Partial Reprogramming. Cell 167, 1719–1733.e12 (2016).

75. B. D. Hale, Y. Severin, F. Graebnitz, D. Stark, D. Guignard, J. Mena, Y. Festl, S. Lee, J. Hanimann, N. S. Zangger, M. Meier, D. Goslings, O. Lamprecht, B. M. Frey, A. Oxenius, B. Snijder, Cellular architecture shapes the naïve T cell response. Science 384, eadh8697 (2024).

76. C. M. Denais, R. M. Gilbert, P. Isermann, A. L. McGregor, M. te Lindert, B. Weigelin, P. M. Davidson, P. Friedl, K. Wolf, J. Lammerding, Nuclear envelope rupture and repair during cancer cell migration. Science 352, 353–358 (2016).

77. Y. Wang, M. Eddison, G. Fleishman, M. Weigert, S. Xu, T. Wang, K. Rokicki, C. Goina, F. E. Henry, A. L. Lemire, U. Schmidt, H. Yang, K. Svoboda, E. W. Myers, S. Saalfeld, W. Korff, S. M. Sternson, P. W. Tillberg, EASI-FISH for thick tissue defines lateral hypothalamus spatio-molecular organization. Cell 184, 6361–6377.e24 (2021).

78. Y. Bai, B. Zhu, J.-P. Oliveria, B. J. Cannon, D. Feyaerts, M. Bosse, K. Vijayaragavan, N. F. Greenwald, D. Phillips, C. M. Schürch, S. M. Naik, E. A. Ganio, B. Gaudilliere, S. J. Rodig, M. B. Miller, M. Angelo, S. C. Bendall, X. Rovira-Clavé, G. P. Nolan, S. Jiang, Expanded vacuum-stable gels for multiplexed high-resolution spatial histopathology. Nat. Commun. 14, 4013 (2023).

79. J.-B. Chang, F. Chen, Y.-G. Yoon, E. E. Jung, H. Babcock, J. S. Kang, S. Asano, H.-J. Suk, N. Pak, P. W. Tillberg, A. T. Wassie, D. Cai, E. S. Boyden, Iterative expansion microscopy. Nat. Methods 14, 593–599 (2017).

80. O. M’Saad, J. Bewersdorf, Light microscopy of proteins in their ultrastructural context. Nat. Commun. 11, 3850 (2020).

81. C. Condylis, A. Ghanbari, N. Manjrekar, K. Bistrong, S. Yao, Z. Yao, T. N. Nguyen, H. Zeng, B. Tasic, J. L. Chen, Dense functional and molecular readout of a circuit hub in sensory cortex. Science 375, eabl5981 (2022).

82. Q. Li, Z. Lin, R. Liu, X. Tang, J. Huang, Y. He, X. Sui, W. Tian, H. Shen, H. Zhou, H. Sheng, H. Shi, L. Xiao, X. Wang, J. Liu, Multimodal charting of molecular and functional cell states via in situ electro-sequencing. Cell 186, 2002–2017.e21 (2023).

83. T. J. Chozinski, A. R. Halpern, H. Okawa, H.-J. Kim, G. J. Tremel, R. O. L. Wong, J. C. Vaughan, Expansion microscopy with conventional antibodies and fluorescent proteins. Nat. Methods 13, 485–488 (2016).

84. S. Chen, Y. Zhou, Y. Chen, J. Gu, fastp: an ultra-fast all-in-one FASTQ preprocessor. Bioinformatics 34, i884–i890 (2018).

85. B. Langmead, S. L. Salzberg, Fast gapped-read alignment with Bowtie 2. Nat. Methods 9, 357–359 (2012).

86. H. Li, B. Handsaker, A. Wysoker, T. Fennell, J. Ruan, N. Homer, G. Marth, G. Abecasis, R. Durbin, 1000 Genome Project Data Processing Subgroup, The Sequence Alignment/Map format and SAMtools. Bioinformatics 25, 2078 (2009).

87. T. Smith, A. Heger, I. Sudbery, UMI-tools: modeling sequencing errors in Unique Molecular Identifiers to improve quantification accuracy. Genome Res. 27, 491 (2017).

88. W. Bao, K. K. Kojima, O. Kohany, Repbase Update, a database of repetitive elements in eukaryotic genomes. Mob. DNA 6 (2015).

89. A. R. Quinlan, I. M. Hall, BEDTools: a flexible suite of utilities for comparing genomic features. Bioinformatics 26, 841–842 (2010).

91. A. S. Berg, D. Kutra, T. Kroeger, C. N. Straehle, B. X. Kausler, C. Haubold, M. Schiegg, J. Ales, T. Beier, M. Rudy, K. Eren, J. I. Cervantes, B. Xu, F. Beuttenmueller, A. Wolny, C. Zhang, U. Koethe, F. A. Hamprecht, A. Kreshuk, ilastik: interactive machine learning for (bio)image analysis. Nat. Methods 16, 1226–1232 (2019).

90. S. S. Rao, M. H. Huntley, N. C. Durand, E. K. Stamenova, I. D. Bochkov, J. T. Robinson, A. L. Sanborn, I. Machol, A. D. Omer, E. S. Lander, E. L. Aiden, A 3D map of the human genome at kilobase resolution reveals principles of chromatin looping. Cell 159 (2014).

91. X. Wang, F. Yue, HiCLift: a fast and efficient tool for converting chromatin interaction data between genome assemblies. Bioinformatics 39, btad389 (2023).

92. N. C. Durand, M. S. Shamim, I. Machol, S. S. P. Rao, M. H. Huntley, E. S. Lander, E. L. Aiden, Juicer provides a one-click system for analyzing loop-resolution Hi-C experiments. Cell systems 3, 95 (2016).

93. ENCODE Project Consortium, An integrated encyclopedia of DNA elements in the human genome. Nature 489, 57–74 (2012).

